# Symbiotic and commensal root microbes are key determinants of plant responses to rising CO_2_

**DOI:** 10.64898/2026.11.07.764780

**Authors:** Marina Cotta, Paul Kajdewicz, Lysandros Anastasopoulos, Carlos Duran-Ballesteros, Jianbei Huang, Karl Kübler, Martin Göbel, Susan Trumbore, Paul Schulze-Lefert, Ruben Garrido-Oter

## Abstract

Rising atmospheric CO_2_ stimulates plant growth, but this effect is often constrained by nutrient availability. While beneficial soil-derived microbial symbionts such as nitrogen-fixing rhizobia and mycorrhizal fungi are known to mitigate this limitation, the role of plant commensal microbiota remains unclear. Here, using a gnotobiotic growth system and the model legume *Lotus japonicus*, we show that the magnitude of plant responses to elevated CO_2_ is largely explained by the composition of its root-associated microbial community. Axenic plants exhibited limited physiological responses to increasing CO_2_ but inoculation with symbiotic and commensal microbes amplified this effect. Commensal bacteria alone, while not increasing overall biomass, shifted plant carbon allocation belowground. Microbial abundances in roots increased with atmospheric CO_2_, even when normalized to plant DNA. The strongest plant and microbial responses were observed when rhizobia, arbuscular mycorrhizal fungi, and the commensal bacterial community were all present. These findings indicate that multi-partner microbial associations will shape both plant growth and the allocation of carbon belowground as atmospheric CO_2_ concentrations continue to rise.

## Introduction

The rising concentration of atmospheric CO_2_ is altering the functioning of terrestrial ecosystems and soil carbon cycles at a global scale *(1–4)*. Elevated CO_2_ is known to stimulate plant growth and increase productivity, a phenomenon which is known as the CO_2_ fertilization effect, and which has long been recognized as a key mechanism by which land vegetation acts as a major carbon sink (5–7). However, the magnitude of this effect is constrained by nutrient availability, particularly nitrogen (N) and phosphorus (P), which are essential mineral macronutrients for producing biomass and, consequently, carbon sequestration. Nutrient limitation can reduce or even negate the CO_2_ fertilization effect *(8, 9)*, posing uncertainty in global carbon cycle models, with major implications for global carbon budget predictions *(10)*.

This macronutrient limitation can be alleviated by associations with symbiotic microbial partners, notably those involving arbuscular mycorrhizal fungi (AMF) and nitrogen-fixing rhizobia. Associations with AMF are widespread in land plants and result in increased P acquisition, while rhizobia induce the formation of root nodules in legumes, where the bacterial symbiont fixes atmospheric N and provides it to its host in the form of ammonia. The effectiveness of these interactions, and how they modify the impact of increased CO_2_ levels on host growth varies with the plant nutritional status and with the symbiosis functional type. For example, ectomycorrhizal (ECM) trees typically show strong CO_2_ fertilization effects, even under low N, while AMF-associated plants often remain N-limited *(11)*. Nodule symbioses in legumes can sustain a stronger growth response to elevated CO_2_ by supplying fixed nitrogen *(12–15)*, although the stimulation of fixation varies with legume species and soil nitrogen, and can fade over years *(16, 17)*. These findings establish microbial functional types as critical mediators of plant responses to rising CO_2_ concentrations, with potential impact on terrestrial carbon cycling.

However, while the role of nutrient limitation and its alleviation by AM fungi and N-fixing rhizobia are well characterized, the influence of the taxonomically diverse community of commensal microbes that colonize the root and rhizosphere, known as the root microbiota, remains poorly understood. Such interactions could influence not only plant growth, but also patterns of root exudation, rhizosphere colonization, and soil carbon fluxes *(18)*, with cascading implications for nutrient cycling and long-term carbon storage *(19)*. In addition, microbial uptake of plant-derived carbon increases microbial respiration, with variable effects on microbial biomass, potentially tipping the balance between soil carbon sequestration and loss *(10, 20–23)*. Furthermore, the physiological responses and growth dynamics of soil microbes are themselves shaped by changing CO_2_ conditions, adding an additional layer of complexity *(24)*. Deciphering how these microbial responses are organized across functional groups remains a major challenge for global change biology. Experiments under elevated CO_2_ have so far compared plants with and without a single symbiont in sterilized substrate *(13, 15).* However, whether a taxonomically complex root microbiota alters the response of a plant to rising CO_2_ has never been tested against germ-free plants.

To address this, we implemented a reductionist approach to isolate microbial contributions to plant carbon dynamics under varying CO_2_. Using a gnotobiotic system with controlled environmental conditions and automated gas exchange monitoring, we cultivated the model legume *Lotus japonicus* under different atmospheric CO_2_ concentrations. This system enabled us to disentangle the roles of three microbial inputs: AM fungi, nitrogen-fixing rhizobia, and a defined synthetic bacterial community of root commensals, alone and in combination, against germ-free plants. We hypothesized that rising CO_2_ alters the balance between photosynthesis and respiration, and that microbial partners modulate plant responses by affecting nutrient availability and carbon allocation between source and sink tissues. Our aim was to quantify how single and combined microbial associations shape plant growth, carbon fluxes, and microbial colonization along a CO_2_ gradient. We show that the magnitude of the plant response to elevated CO_2_ depends on the composition of its root microbiota, and that the strongest responses were observed when rhizobia, mycorrhizal fungi, and commensal bacteria were all present.

## Results

### Areductionist system for gnotobiotic plant growth under varying CO_2_ concentrations

To understand how changes in atmospheric CO_2_ influence plant physiology and root microbiota assembly, we required a system that combined precise environmental control with the capacity to manipulate microbial composition. Natural soils and field experiments offer limited control over abiotic variables and microbial inputs, making it difficult to isolate the specific impacts of varying CO_2_ concentrations. To overcome these limitations, we developed a custom gnotobiotic growth platform, the EcoChamber, which allows plants to grow in sterile peat substrate under controlled humidity, temperature, light, and defined CO_2_ levels, while being colonized by synthetic microbial communities (SynComs) of known composition.

The EcoChamber system is built around sealed plexiglass enclosures that house plastic pots filled with a sterilized peat substrate that can be individually inoculated with defined microbial SynComs (see materials and methods). Germ-free *L. japonicus* seedlings are then planted into the inoculated substrate, ensuring that microbial colonization originates only from the defined inputs. Each chamber contains 20 pots, all receiving the same microbial treatment to prevent microbial cross-contamination. SynComs were derived from a previously established *L. japonicus* root and nodule culture collection (*25; Lj*-SPHERE), and include taxonomically diverse commensal bacteria representing 15 bacterial families spanning the four dominant phyla of the root microbiota (**Table S1**). In addition to the SynCom, plants can also be inoculated with nodule-forming, nitrogen-fixing rhizobia and/or AMF symbionts. The chambers are connected to automated air and water circulation systems (**Fig. 1A**), and placed inside climate-controlled growth cabinets where light intensity, humidity, and temperature can be adjusted. These cabinets are also equipped with CO_2_ scrubbers and connected to gas inputs (**Fig. 1B**), enabling stable maintenance of CO_2_ concentrations at defined levels.

**Figure 1.**
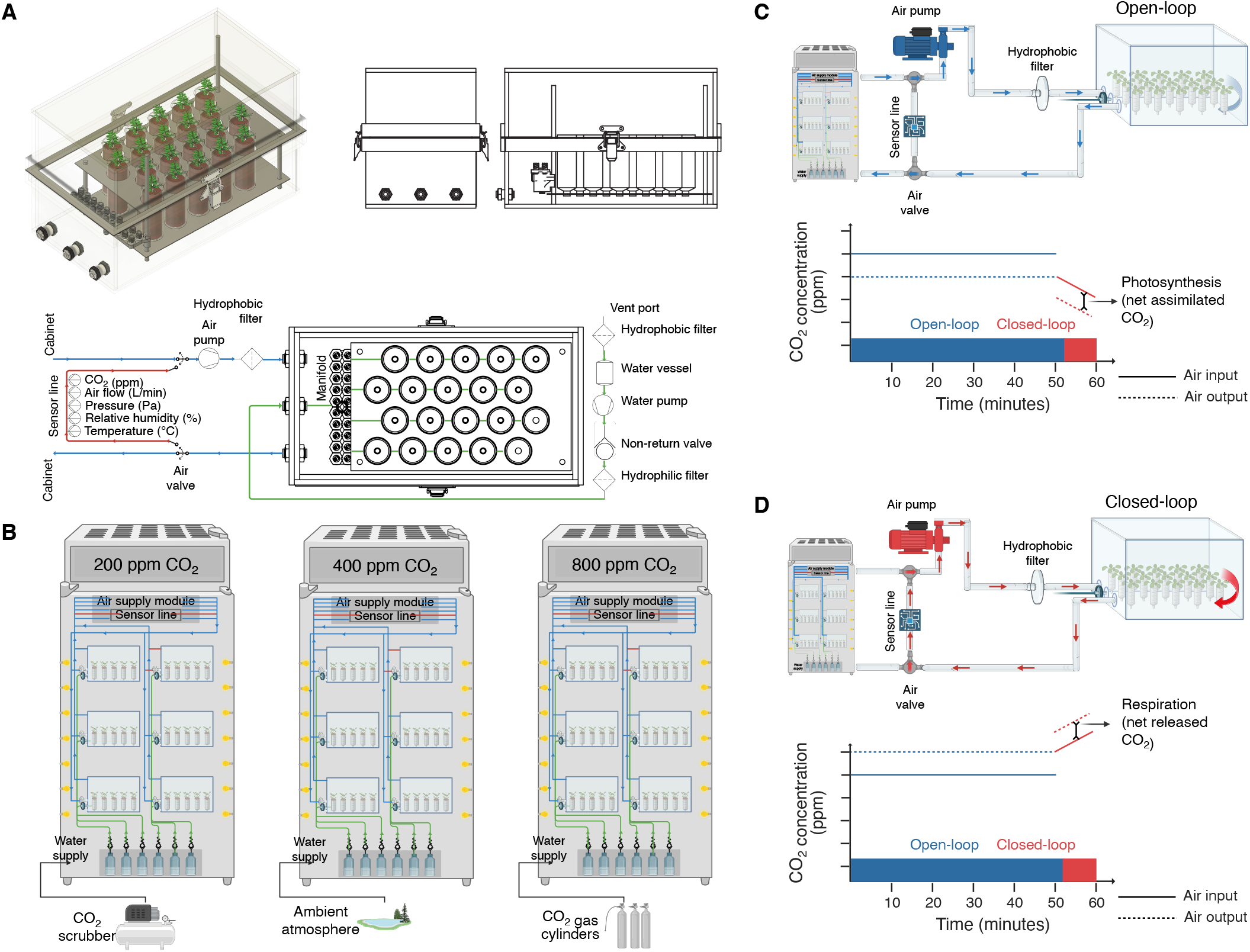
Design and operation of the EcoChamber platform. **(A)** The EcoChamber growth system. Top, isometric and orthographic views of one unit holding 20 FlowPots. Bottom, air and water circuits of a unit. Cabinet air enters through an air pump and a hydrophobic filter and passes a sensor line that records CO_2_ concentration, air flow, pressure, relative humidity, and temperature before returning to the cabinet. Sterile water reaches the pots through a vent port, a water vessel, a water pump, a non-return valve, and a hydrophilic filter. **(B)** Six EcoChamber units inside each of three growth cabinets with controlled light, temperature, and humidity, held at 200 ppm (CO_2_ scrubber), 400 ppm (ambient atmosphere), and 800 ppm CO_2_ (gas cylinders). An air supply module and a sensor line serve the six units of a cabinet. Each unit operates independently, allowing full-factorial combinations of microbial treatments and atmospheric CO_2_ regimes. **(C-D)** Open-loop **(C)** and closed-loop **(D)** airflow configurations, each with the CO_2_ concentration of the air entering (solid line) and leaving (dotted line) the unit over one 60-minute cycle. In open-loop mode, cabinet air flows through the unit. In closed-loop mode, the air valve recirculates the air of the unit through the sensor line, and the CO_2_ concentration falls when net assimilation exceeds respiration **(C)** or rises when net respiration exceeds assimilation **(D)**. The slope of the concentration over the closed-loop interval gives the net CO_2_ flux of the unit.

Under standard operation, air from the growth cabinet circulates through each EcoChamber in an open-loop configuration (**Fig. 1C**), enabling continuous supply of CO_2_. To measure plant-driven changes in CO_2_ concentrations, the system can switch to a closed-loop operation mode (**Fig. 1D**), where air from each chamber recirculates within a sealed circuit and passes through a sensor array that monitors humidity, pressure, and CO_2_ levels in real time. During these closed cycles, measured changes in CO_2_ concentrations reflect the balance of photosynthetic uptake and respiratory release. Positive slopes in CO_2_ concentrations, typically observed during the night, correspond to net respiration, which includes contributions from plant shoots, roots, and microbial activity in the soil. Conversely, negative slopes, observed during the day period, are the consequence of net assimilation by the plant canopy. A limitation of the system is that it does not separate above- and belowground compartments, so net CO_2_ fluxes reflect photosynthetic assimilation in the shoot combined with ongoing root and microbial respiration in soil. Each growth cabinet accommodates up to six EcoChambers, which are connected to the sensor array via automated valves. For 9 minutes every hour, air from each chamber circulates through the sensor array in a closed-loop mode, enabling high-resolution monitoring of CO_2_ fluxes across all chambers. All system operations, data logging, and flux calculations are automated and managed by custom software (see materials and methods).

Three independent growth cabinets are used in parallel (**Fig. 1B**), each configured with identical light, temperature, and humidity settings, but maintained at one of three distinct CO_2_ levels: 200 ppm, representing pre-industrial conditions; 400 ppm, approximating current ambient levels; and 800 ppm, simulating projected future atmospheric concentrations by the end of the century *(26)*. Each cabinet contains up to six gnotobiotic chambers, enabling parallel testing of microbial treatments across CO_2_ regimes. The EcoChamber platform thus provides a controlled environment to quantify gas exchange dynamics and plant-microbe interactions under shifting CO_2_ regimes.

### CO_2_ fertilization effect on L. japonicus depends on microbial associations

To test how root-associated microbes modulate plant responses to rising atmospheric CO_2_, we grew *L. japonicus* in EcoChambers using three CO_2_ concentrations (200, 400, and 800 ppm), in the presence or absence of microbial partners for five weeks. We selected *L. japonicus* for its capacity to form functional associations with both AMF (*Rhizophagus irregularis*) and nitrogen-fixing rhizobia (*Mesorhizobium loti*), and for the availability of a collection of well-characterized root- and nodule-derived bacterial cultures (25). This allowed us to design a modular gnotobiotic experiment combining AMF, *M. loti*, and a bacterial SynCom composed of 15 diverse isolates derived from natural root communities.

Under axenic conditions, we observed a small but significant CO_2_ fertilization effect across nearly all plant traits measured, including shoot and root fresh weight, shoot length, chlorophyll content, and total plant weight (**Fig. 2A-E**). Compared to ambient levels (400 ppm), shoot and root fresh weight increased by 37% and 29%, respectively, under elevated CO_2_ (800 ppm), and decreased by 49% and 86% under low CO_2_ (200 ppm; **Fig. 2A, D**). These results confirm that the EcoChamber system captures expected CO_2_-driven plant physiological responses and provides a baseline against which the effects of different microbial associations can be compared (dashed lines in **Fig. 2**).

**Figure 2.**
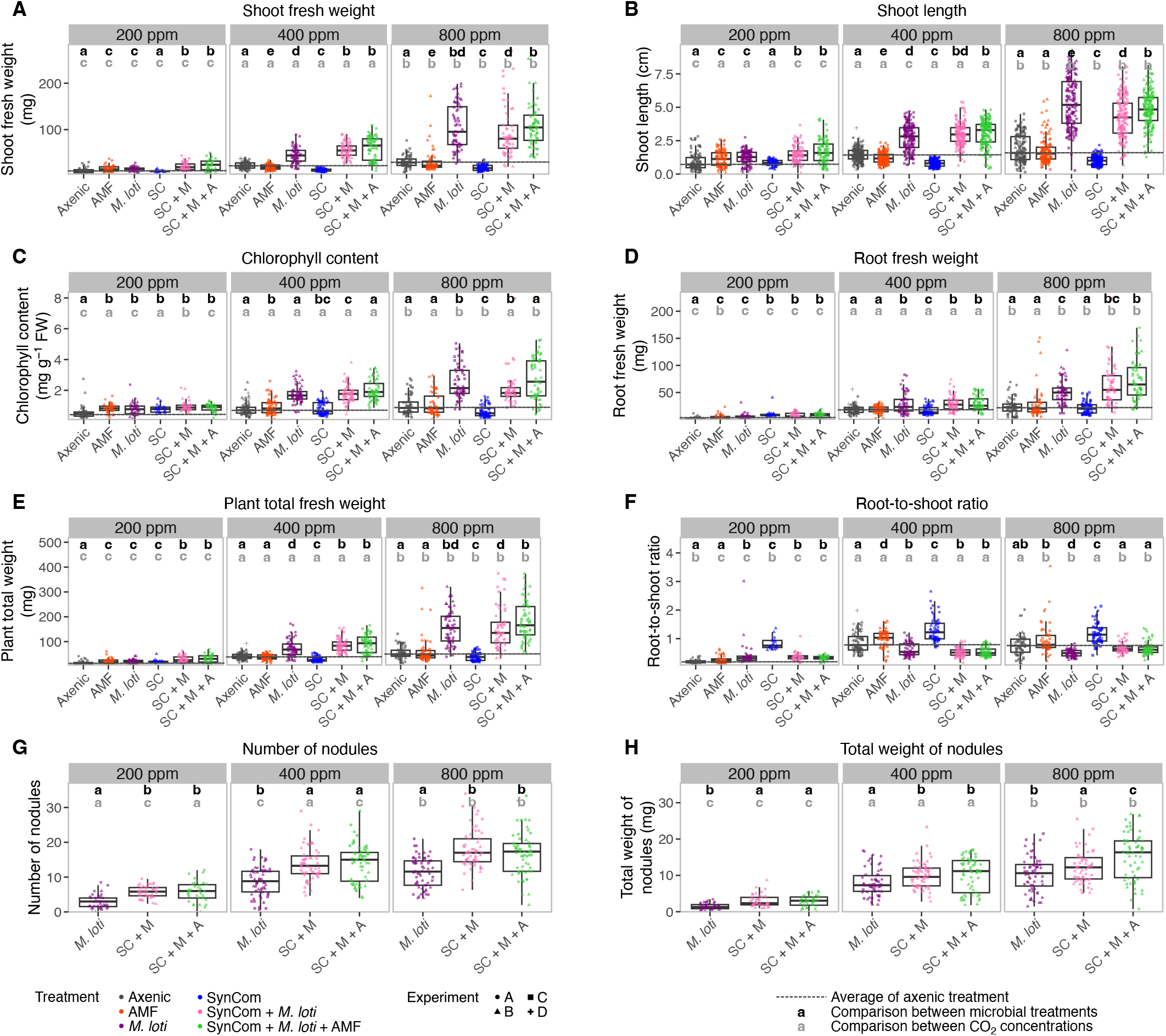
CO_2_ fertilization effect on *L. japonicus* treated with different root-associated microbial communities. *L. japonicus* plants were grown for five weeks under six microbial treatments and three atmospheric CO_2_ concentrations. **(A)** Shoot fresh weight, **(B)** shoot length, **(C)** chlorophyll content, **(D)** root fresh weight, (E) plant total fresh weight, **(F)** root-to-shoot ratio, **(G)** number of nodules, and (H) total weight of nodules. Nodule traits are shown for the treatments containing *M. loti*. Boxes show the median and interquartile range, whiskers extend to 1.5 times the interquartile range, and the dashed line marks the mean of the axenic treatment at each CO_2_ level. Each data point is the mean of the plants in one pot (mostly 3), except in **(B)**, where each plant was measured separately. Data are pooled from two to four independent experiments per treatment and CO_2_ level, with 20 pots per treatment and CO_2_ level in each experiment, and symbols denote the experiment (A to D). Black letters indicate differences between microbial treatments within a CO_2_ level and grey letters differences between CO_2_ levels within a treatment. Statistical significance was determined by Kruskal-Wallis or ANOVA followed by post hoc tests (see materials and methods). Treatment abbreviations: SC, SynCom; SC + M, SynCom + *M. loti*; SC + M + A, SynCom + *M. loti* + AMF.

Under ambient atmospheric conditions (400 ppm), inoculation with *M. loti* led to significant improvements in all plant performance parameters relative to axenic controls (**Fig. 2A-E**). Further increases were observed when *M. loti* was combined with the bacterial SynCom, or when all three microbial treatments were present, including AMF. By contrast, treatments with AMF or SynCom alone did not improve plant growth. The former result was expected as the peat substrate used in these experiments is not limited in bioavailable P (Table S2) and thus offsets the beneficial effect of AMF on plant growth. The absence of a growth benefit from the SynCom alone contrasts with the growth promotion by Lj-SPHERE SynComs observed previously in the presence of 1 mM nitrate *(25)*, and is consistent with nitrogen being the limiting nutrient in our substrate. Although AMF inoculation did not enhance growth, root colonization by *R. irregularis* was detected across all CO_2_ levels using qPCR targeting a region of the fungal mitochondrial DNA (**Fig. S1**), indicating that AMF colonization alone was insufficient to boost plant growth in the experimental conditions tested.

When plants were grown under elevated CO_2_, the effect of microbial inoculation was markedly increased. *Plants* inoculated with *M. loti* alone showed a 137% increase in shoot fresh weight under 800 ppm compared to 400 ppm (**Fig. 2A**). *Plants* treated with *M. loti* plus SynCom, and those combining all three microbial treatments, showed increases of 73% and 82%, respectively, when comparing 800 ppm to 400 ppm conditions. These increases exceeded the gains observed in axenic plants under elevated CO_2_, indicating a synergistic interaction between microbial treatment and plant physiological responses. Across all traits measured, the highest average values for shoot biomass, chlorophyll content, and root biomass were observed when *M. loti*, AMF, and the commensal SynCom were present together (**Fig. 2A, C, D**), although not all differences were statistically significant. This suggests that the presence of multiple microbial partners can produce additive or synergistic effects that enhance plant performance beyond what is achieved by associations with either AMF or rhizobial symbionts alone.

As expected, nodulation was strictly dependent on the presence of *M. loti*. No nodules were found in axenic, AMF-only, or SynCom-only treatments under any CO_2_ condition (**Fig. 2G-H**). Among *M. loti*-containing treatments, the number and total weight of nodules increased with rising CO_2_ levels, consistent with enhanced symbiotic nitrogen fixation under enriched carbon conditions. Notably, plants co-inoculated with *M. loti* and the bacterial SynCom, with or without AMF, showed significantly higher nodulation than those treated with *M. loti* alone (**Fig. 2G**). The available data, however, are insufficient to determine if this increased nodulation is the consequence of indirect effects of the bacterial SynCom on the plant metabolic status or the result of direct microbial interactions with *M. loti* or with the plant symbiotic signaling pathway.

Unlike other growth traits, root-to-shoot ratios exhibited an unanticipated and markedly distinct response pattern (**Fig. 2F**). The highest values were observed in plants inoculated with the SynCom alone, followed by AMF-only, and axenic treatments. In contrast, plants associated with *M. loti* showed lower root-to-shoot ratios at 400 and 800 ppm, consistent with greater investment in shoot biomass when nitrogen is readily available. The higher ratios observed in the SynCom-only treatment, despite limited impact on total plant biomass, suggest a commensal influence on plant resource allocation. Whether this reflects a stress-induced shift that selectively inhibits above-ground plant growth or an adaptive physiological response to microbial signals that promote below-ground organ growth remains unclear. However, such microbe-driven effects on biomass allocation have potential implications for belowground carbon deposition under rising atmospheric CO_2_.

Taken together, these findings show that microbial associations not only influence the magnitude of the plant CO_2_ fertilization response but also alter carbon allocation along the root-shoot axis. In particular, the full benefit of CO_2_ fertilization in *L. japonicus* required its nitrogen-fixing symbiont, and was further enhanced when *M. loti* was combined with root-derived commensal communities and AMF. As global CO_2_ levels rise, such interactions are likely to affect not only plant growth and productivity but also have implications for soil carbon and nitrogen fluxes in both managed and natural ecosystems.

### Symbiotic and commensal microbes shape carbon assimilation responses to rising CO_2_

To determine whether microbial associations modulate not only plant growth but also net carbon fluxes, we used the EcoChamber system to follow CO_2_ dynamics during approximately 35 days of *L. japonicus* growth under the three previously described atmospheric CO_2_ concentrations (**Fig. 3A-C**). To estimate gas fluxes, each EcoChamber was periodically switched to a closed-loop configuration allowing measurement of CO_2_ concentrations with an accuracy of ±3 ppm, sampled once per second (**Fig. 1A**). During each closed-loop cycle, changes in CO_2_ concentration were recorded and fitted with a regression model (**Fig. 1C**; see materials and methods). The slope of the curve corresponds to the net CO_2_ flux in the chamber. Negative slopes reflect net CO_2_ assimilation, and positive slopes indicate net CO_2_ release. As both photosynthesis and respiration occur simultaneously, and we do not separate above- and belowground compartments, measured values reflect the net balance of shoot assimilation minus respiration by shoots, roots, and soil microbes. These values are referred to as net fluxes throughout.

**Figure 3.**
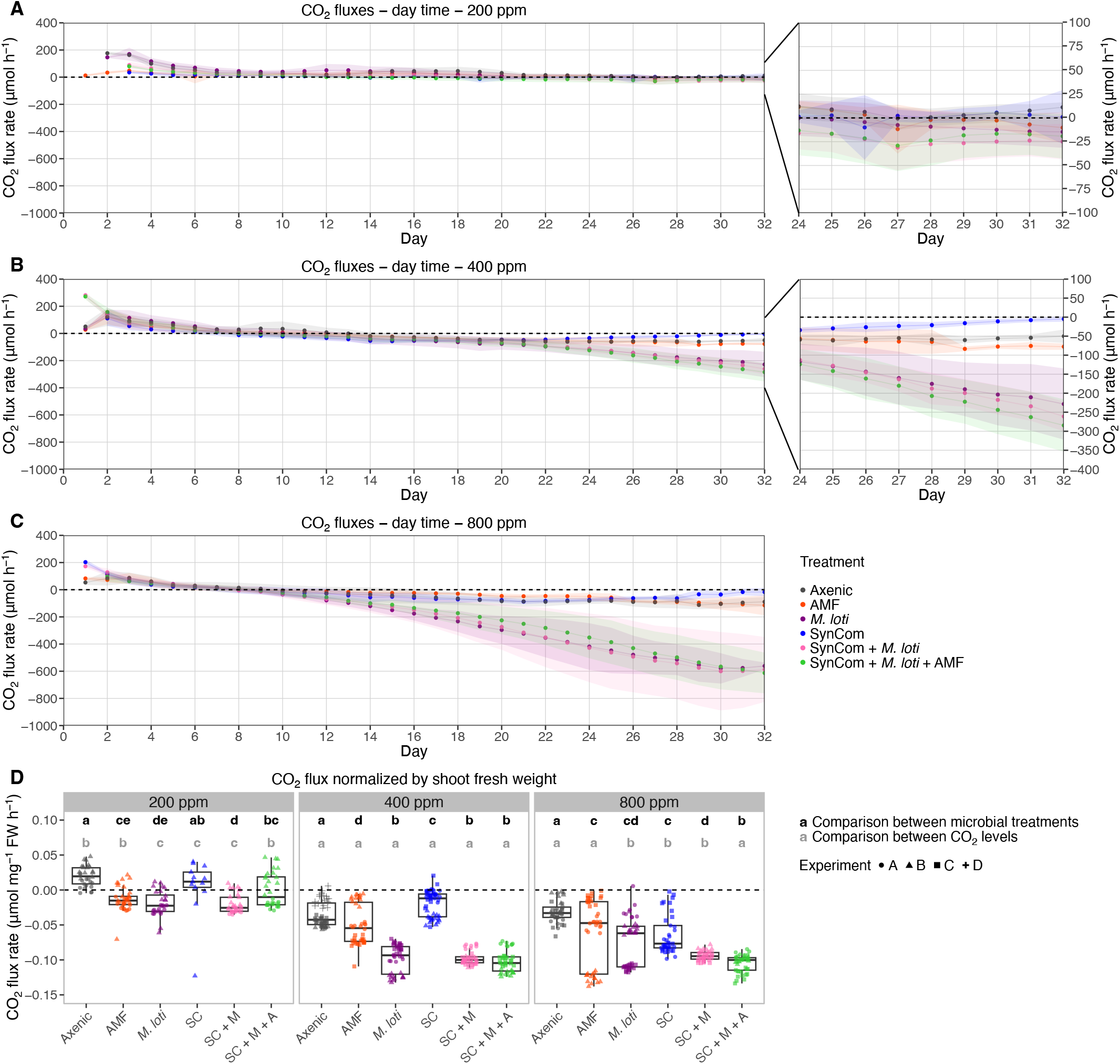
Net CO_2_ fluxes during daytime across CO_2_ and microbial treatments. CO_2_ fluxes were measured over approximately 35 days of *L. japonicus* growth under 200 ppm **(A)**, 400 ppm **(B)**, and 800 ppm **(C)** atmospheric CO_2_. Data points are daily means of the hourly daytime flux measurements of individual chambers, from two to four independent experiments, and shaded bands show the standard deviation of each treatment. The insets in **(A)** and **(B)** show days 24 to 32 at an expanded scale. **(D)** Final-day CO_2_ flux measurements normalized by shoot fresh weight. Each data point is one hourly flux measurement during the daytime of the final day (10 per chamber), from two to four independent experiments, with symbols denoting the experiment (A to D). Positive values reflect net respiration, while negative values reflect net CO_2_ assimilation. Black letters indicate differences between microbial treatments within a CO_2_ level and grey letters differences between CO_2_ levels within a treatment. Statistical significance was determined by Kruskal-Wallis or ANOVA. Treatment abbreviations: SC, SynCom; SC + M, SynCom + *M. loti*; SC + M + A, SynCom + *M. loti* + AMF.

Under ambient CO_2_ conditions (400 ppm; **Fig. 3B**), fluxes remained positive during the day for all treatments early in the experiment, indicating that respiratory outputs exceeded assimilation. This process is likely attributable to the mobilization of seedborne carbon resources by the plant, combined with microbial respiration in the soil substrate. From day 12 onwards, net fluxes became negative for most treatments, marking the onset of net daytime carbon uptake. Starting at day 24, and until the end of the experiment, treatments including *M. loti*, either alone, with SynCom, or with both SynCom and AMF, exhibited significantly larger negative net fluxes on average (−230 to −331 μmol h^−1^) than axenic controls, SynCom-only, or AMF-only conditions (−41, −12, −58 μmol h^−1^ on average, respectively). These results are consistent with increased nitrogen availability supporting greater plant biomass and photosynthetic capacity, and mirror the shoot fresh weight differences observed at harvest (**Fig. 2A**).

At low CO_2_ (200 ppm, **Fig. 3A**), daytime fluxes remained positive for longer, and net negative fluxes only appeared later in the experiment for all treatments (days 27 to 32), except for axenic and SynCom-only conditions. Overall, at 200 ppm net flux magnitudes were smaller than those observed at 400 ppm, and only chambers with *M. loti* inoculated plants showed significantly different patterns compared to axenic controls. When comparing ambient CO_2_ and elevated CO_2_ conditions (400 and 800 ppm, respectively), net fluxes not only turned negative earlier in the experiment (from day 12 compared to day 14 for axenic conditions) but also reached greater magnitudes overall (**Fig. 3B-C**). Treatments containing *M. loti* again showed the strongest negative fluxes, ranging from −534 to −627 μmol h^−1^, approximately 13-fold greater than axenic controls. While fluxes for AMF-only were similar to axenic conditions, SynCom-only chambers displayed significantly weaker net assimilation (−27 μmol h^−1^), suggesting either increased respiration, decreased assimilation, or both.

To determine whether differences in gas fluxes could be attributed solely to differences in aboveground plant biomass, we normalized final-day fluxes by shoot fresh weight at harvest (**Fig. 3D**). This analysis revealed that under ambient conditions (400 ppm), all microbial treatments showed stronger CO_2_ assimilation per gram of plant shoot biomass compared to their counterparts under 200 ppm, suggesting increased normalized photosynthetic performance. However, no treatment showed higher normalized assimilation at 800 ppm than at 400 ppm, except for SynCom-only, which exhibited a significantly stronger negative flux per unit biomass under elevated CO_2_ (**Fig. 3D**). Surprisingly, despite having lower shoot biomass than axenic or symbiont-inoculated plants (**Fig. 2A**), SynCom-only plants achieved a higher normalized CO_2_ drawdown rate than axenic plants (**Fig. 3D**).

Several explanations could account for this result. It is possible that the bacterial SynCom enhances the photosynthetic capacity of *L. japonicus* per unit of leaf tissue, for instance by modulating leaf growth traits. Alternatively, the presence of bacterial commensals might increase belowground carbon sink strength, stimulating photosynthesis via altered source-sink dynamics. A third possibility is that SynCom-only treatments exhibit reduced microbial respiration, resulting in a more negative net flux. These hypotheses are not mutually exclusive, and further physiological and molecular analyses will be needed to disentangle their relative contributions.

In sum, these findings indicate that microbial associations shape not only the magnitude of differences in plant growth under changing CO_2_ levels, but also the trajectory of net carbon fluxes over time. The presence of symbiotic rhizobia accelerates both the onset and magnitude of net carbon gain, particularly under elevated CO_2_, while commensal microbes modulate carbon assimilation in more subtle or indirect ways. Together, these results highlight a complex and dynamic interaction between microbial association type, atmospheric CO_2_ concentration, and plant source-sink carbon balance.

### Atmospheric CO_2_ levels shape the composition and abundance of the root microbiota

To determine how rising atmospheric CO_2_ concentrations affect the composition of the root microbiota, we performed bacterial 16S rRNA gene amplicon profiling and absolute quantification of bacterial abundance from *L. japonicus* plants grown under the same conditions described above. Our analyses focused on both the root (root surface and interior) and rhizosphere (surrounding peat substrate) compartments, and included comparisons across microbial treatments and CO_2_ concentrations.

Principal Coordinate Analysis (PCoA) of amplicon profiles revealed a clear and consistent separation between microbial communities in the root and rhizosphere, after in silico depletion of reads belonging to *M. loti* (**Fig. 4A**). This separation by compartment was evident across all CO_2_ conditions and independent of the microbial treatment applied, confirming that the plant exerts strong spatial filtering over the SynCom inoculated in the growth substrate.

**Figure 4.**
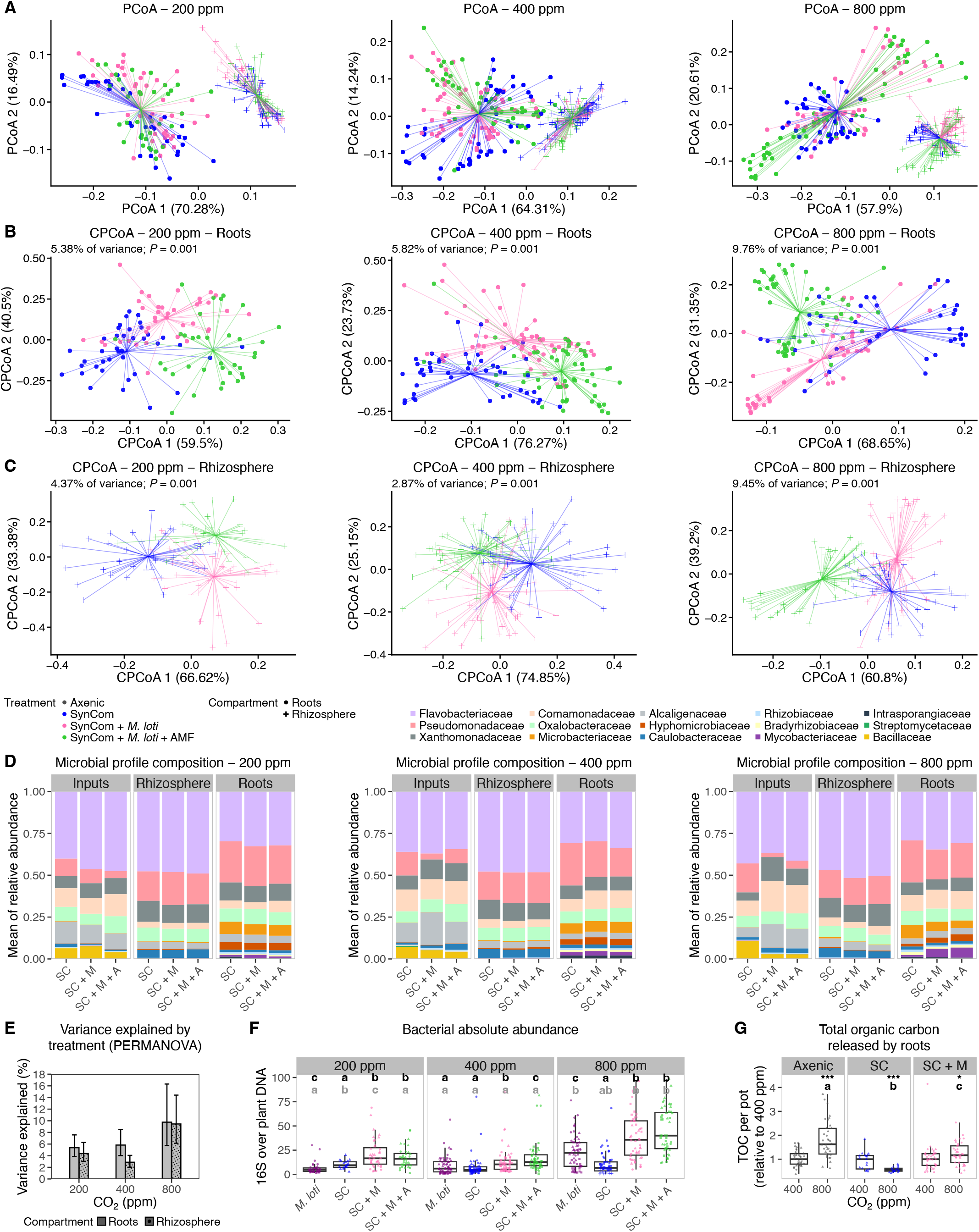
CO_2_-dependent effects of microbial treatment on root- and rhizosphere-associated communities. **(A)** Principal Coordinate Analysis (PCoA) of Bray-Curtis dissimilarities between 16S rRNA profiles from samples of root and rhizosphere compartments across all CO_2_ concentrations. PCoA of bacterial community profiles constrained by microbial treatment at each CO_2_ level from root **(B)** or rhizosphere **(C)** samples. **(D)** Compositional profiles of SynCom inputs and communities retrieved from roots and rhizosphere. **(E)** PERMANOVA analysis of treatment effects on community composition, showing increased variance explained at elevated CO_2_. **(F)** Bacterial abundance in root systems quantified by qPCR of 16S rRNA normalized to the *L. japonicus Nfr5* gene. Each data point reflects qPCR results from the pooled plants of one pot (mostly 3), from two to four independent experiments per treatment and CO_2_ level, with 20 pots per treatment and CO_2_ level in each experiment, and symbols denote the experiment (A to D). Black letters indicate differences between microbial treatments within a CO_2_ level and grey letters differences between CO_2_ levels within a treatment. Statistical significance was determined by Kruskal-Wallis or ANOVA. Treatment abbreviations: SC, SynCom; SC + M, SynCom + *M. loti*; SC + M + A, SynCom + *M. loti* + AMF. **(G)** Root-derived organic carbon released over 24 h into sterile water by the root systems of axenic, SynCom, and SynCom + *M. loti* plants grown at 400 or 800 ppm, in two independent experiments (triangles and circles). The SynCom treatment was tested in one experiment. Each data point represents one pot. Values are per pot and normalized to the median of the 400 ppm pots of the same treatment and experiment (*n* = 19 to 40 pots per group). Asterisks mark differences between 800 and 400 ppm within each treatment (Wilcoxon rank-sum test, *, *P* < 0.05; ***, *P* < 0.001), and letters mark treatments whose CO_2_ responses differ from each other (*P* < 0.001).

To assess how microbial treatments shaped bacterial community structure under each CO_2_ condition, we constrained the principal coordinate analysis by treatment. This revealed significant differences in community composition between treatments (SynCom only, SynCom + *M. loti*, and SynCom + *M. loti* + AMF), with the magnitude of the separation increasing when comparing 400 to 800 ppm (**Fig. 4B, E**). Permutational multivariate analysis of variance (PERMANOVA) showed that microbial treatment explained 5.38%, 5.82%, and 9.76% of the total variance in root community composition at 200, 400, and 800 ppm, respectively (**Fig. 4B, E**). These results suggest that the impact of microbial input on the composition of the root microbiota becomes stronger under elevated CO_2_. A similar analysis for the rhizosphere compartment (**Fig. 4C, E**) also showed treatment-dependent effects, which rose from 4.37% and 2.87% of variance at 200 and 400 ppm to 9.45% at 800 ppm, close to the 9.76% explained in roots at that level. The stronger treatment effect in roots confirms that selection by the plant is a key determinant of root microbiota structure.

Analysis of the taxonomic composition of microbial communities further revealed that root SynComs were characterized by increased relative abundances of strains belonging to the Pseudomonadaceae, Microbacteriaceae, Hyphomicrobiaceae and Mycobacteriaceae families compared to input samples, and decreased relative abundance of members of the Flavobacteriaceae, Comamonadaceae and Alcaligenaceae families (**Fig. 4D**). In contrast, rhizosphere communities showed higher average relative abundance of Flavobacteriaceae compared to roots but a decrease in the abundance of members of the Microbacteriaceae, Hyphomicrobiaceae and Mycobacteriaceae. These compartment-specific differences in taxonomic profiles were consistent across CO_2_ levels, as expected from the analyses of beta-diversity (**Fig. 4A**). However, we also observed smaller but significant differences in the relative abundance of a subset of community members across CO_2_ conditions. The relative abundance of Comamonadaceae and Bradyrhizobiaceae increased with rising CO_2_ levels across all microbial treatments, whereas the relative abundance of Xanthomonadaceae showed the opposite pattern. For some taxa, the correlation with increasing CO_2_ was dependent on the microbial treatment: Mycobacteriaceae showed increased abundance in the presence of microbial symbionts, but decreased abundance on SynCom-only treated plants, while the opposite trend was observed for Rhizobiaceae (**Fig. S2**).

We next asked whether these CO_2_- and treatment-dependent shifts in community composition were accompanied by changes in total microbial load. We quantified bacterial abundance in roots using qPCR of the bacterial 16S rRNA gene normalized to the *L. japonicus* Nfr5 single-copy gene (**Fig. 4F**). Across all CO_2_ conditions, plants inoculated with SynCom + *M. loti* or SynCom + *M. loti* + AMF harbored significantly more bacteria per plant DNA than those treated with SynCom or *M. loti* alone. This result indicates that co-inoculation with commensals and symbionts leads to higher microbial loads in the root.

Bacterial abundances were higher under elevated CO_2_ than under ambient conditions across all microbial treatments (**Fig. 4F**), as previously reported for wheat roots under elevated CO_2_ and high nitrate supply (27). At 800 ppm, the same amount of plant DNA corresponded to significantly higher bacterial DNA, suggesting that higher atmospheric CO_2_ levels increase the capacity of the root system to support microbial colonization. To directly probe whether elevated CO_2_ led to increases in plant carbon release belowground, we quantified the organic carbon released over 24 h by the root systems of axenic, SynCom, and SynCom plus *M. loti* inoculated plants grown at 400 and 800 ppm, in two independent experiments (see materials and methods). Root systems of axenic plants released 1.6-fold more organic carbon at 800 ppm than at 400 ppm, consistently in both experiments (**Fig. 4G**), showing that elevated CO_2_ increases root carbon release in *L. japonicus* in the absence of microbial partners. This response depended on the root microbiota. *Plants* colonized by the SynCom plus *M. loti* showed a smaller increase (1.2-fold), and plants colonized by the SynCom alone released less carbon at 800 than at 400 ppm (0.55-fold), so that the CO_2_ responses of the three treatments differed from each other (**Fig. 4G**). These results show that the increase in root carbon release driven by elevated CO_2_ is reduced in plants colonized by the SynCom plus *M. loti* and abolished in plants colonized by the SynCom alone.

Together, these findings reveal that both the composition and abundance of the root microbiota are affected by atmospheric CO_2_ concentration. While microbial composition becomes increasingly responsive to rising CO_2_ (**Fig. 4E**), the total microbial load also increases (**Fig. 4F**), particularly when commensals and symbionts are combined. Under elevated CO_2_, *M. loti* alone supported higher microbial biomass than SynCom alone, but the highest bacterial loads were found in treatments combining *M. loti* with the SynCom, with or without AMF. These results parallel our earlier observations on increased nodule number and weight under these same combinations (**Fig. 2G-H**), pointing to a functional interaction between symbionts and commensals that becomes more pronounced under elevated CO_2_. Integrating the carbon release measurements with the microbial load data suggests that the root microbiota controls the fate of the additional carbon fixed under elevated CO_2_. In axenic plants, part of the additional assimilate is released from the roots as organic carbon. In plants colonized by the SynCom plus *M. loti*, carbon release rose only modestly while root systems, nodules and bacterial loads increased (**Fig. 2D, G-H, 4F**), suggesting that the additional carbon is retained in the root and by its associated microbes rather than released. Commensals alone, which supply no nitrogen, reduced carbon release under elevated CO_2_ and supported only modest bacterial loads, in line with the limited growth response of these plants (**Fig. 2**). One possibility is that axenic plants invest more carbon in exudation to recruit microbes for nutrient mobilization and protection, a demand that is met once the root is colonized. Alternatively, part of the carbon released by colonized roots may have been taken up by bacteria attached to the root during the 24 h collection, which we did not quantify. Given the strong link between microbial activity and nutrient cycling in soil, these CO_2_-dependent changes in carbon release and microbial biomass have important implications for understanding how plant-microbiota associations will influence soil carbon dynamics in future climate scenarios.

### Microbial community composition shapes multi-trait plant responses to elevated CO_2_

To compare how different microbial associations modulate plant and microbial responses to rising atmospheric CO_2_, we jointly analyzed the major physiological and microbial traits measured (**Fig. 5**, **Fig. S3**). This analysis integrates ten key measurements across CO_2_ concentrations, including plant growth parameters, nodulation patterns, CO_2_ fluxes, and microbial abundance, and provides an overview of how each treatment alters plant and microbial traits in response to increasing CO_2_ levels.

**Figure 5.**
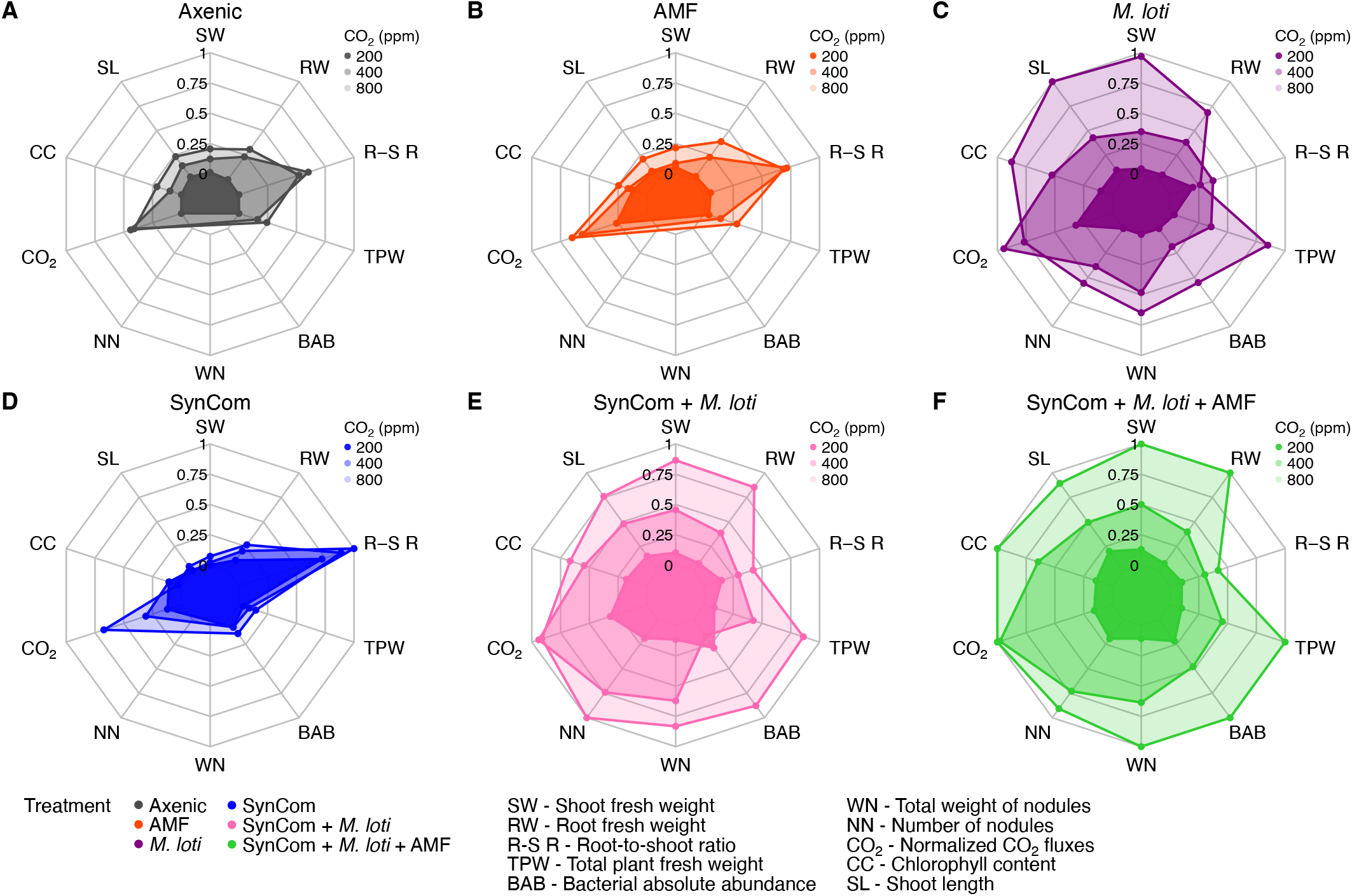
Microbial associations shape multi-trait plant responses to rising CO_2_. Spider plots summarizing the CO_2_-dependent response of *L. japonicus* to different microbial treatments based on ten traits: shoot fresh weight, shoot length, chlorophyll content, root fresh weight, plant total fresh weight, root-to-shoot ratio, nodule number, total nodule weight, bacterial abundance, and normalized CO_2_ fluxes. Each panel corresponds to one microbial treatment: axenic **(A)**, AMF only **(B)**, *M. loti* only **(C)**, SynCom only **(D)**, SynCom + *M. loti* **(E)**, and SynCom + *M. loti* + AMF (F), with the three CO_2_ concentrations shown as nested polygons. Each trait is scaled from 0 (inner ring) to 1 (outer ring). Traits that do not apply to a treatment are absent from its polygon.

In axenic plants, we observed a small but significant CO_2_ fertilization effect when comparing 400 and 800 ppm across most growth-related traits, including shoot fresh weight, shoot length, and chlorophyll content (**Fig. 5A**). However, aside from these biomass gains, the overall shape of the response remains relatively uniform across CO_2_ concentrations, indicating limited physiological plasticity in the absence of microbial partners. AMF-treated plants closely resemble axenic controls (**Fig. 5B**), consistent with previous results and likely explained by the non-limiting amount of phosphorus present in our growth substrate (**Table S2**).

The SynCom-only treatment exhibited a distinct signature (**Fig. 5D**). Across all CO_2_ concentrations, these plants showed minimal changes in growth, nodulation, or chlorophyll content, suggesting that commensals alone cannot support strong responsiveness to elevated CO_2_. However, SynCom-inoculated plants consistently displayed the highest root-to-shoot ratios of any treatment, including under elevated CO_2_. This suggests that interactions with bacterial commensals strongly influence plant decisions on resource allocation to above- or below-ground organs, even in the absence of nitrogen- or phosphorus-providing symbioses. Whether this reflects a developmental response to microbial signaling or an adaptive reallocation of resources under nutrient limitation remains unresolved, but the pattern is remarkably consistent across independent replicates.

By contrast, plants inoculated with *M. loti*, with or without bacterial commensals (**Fig. 5C, E-F**), displayed strong responses to elevated CO_2_ across nearly all traits. In these treatments, gains in shoot and root biomass, chlorophyll content, nodule number and weight, and microbial abundance all increased with rising CO_2_ levels. The combination of *M. loti*, SynCom, and AMF produced the most substantial trait-level gains at 800 ppm across nearly all axes (**Fig. 5F**). In addition, the impact of the SynCom treatment on root-to-shoot ratios was not observed when in combination with *M. loti* and AMF, indicating that functional symbioses can override the altered resource allocation patterns that we observed in the presence of commensal bacterial communities alone. These patterns suggest that the full physiological benefit of elevated CO_2_ requires simultaneous association between the plant host and all three types of microbial partners.

These findings highlight that the root-associated microbiota shape plant CO_2_ responses in a treatment-specific manner. Functional associations with rhizobia and commensals together enhance growth, microbial colonization, and net carbon uptake, particularly under elevated CO_2_, while AMF contribute less to plant performance in the tested conditions, unless paired with other microbial partners. By contrast, under germ-free conditions, the CO_2_ fertilization effect on plant growth is small when CO_2_ concentrations rise. Further, bacterial commensals alone do not enhance growth but consistently shift biomass allocation toward belowground compartments, underscoring their subtle but significant role in altering plant physiology. Taken together, these data support a model in which microbial diversity provides complementary functional benefits that increase under projected future atmospheric CO_2_ conditions.

## Discussion

How plants respond to rising atmospheric CO_2_ is a central question for understanding future terrestrial ecosystems. While the CO_2_ fertilization effect on plant productivity is well established (28), its interaction with the root-associated microbiota remains poorly resolved. Our study reveals that both the magnitude and nature of plant responses to elevated CO_2_ strongly depend on microbial associations, with the strongest responses observed only when commensals, symbiotic rhizobia, and mycorrhizal fungi were all present. Our results, obtained in a controlled gnotobiotic system using the model legume *L. japonicus*, demonstrate that microbial diversity plays a critical role in plant responses to rising atmospheric CO_2_ conditions.

Using the EcoChamber platform, we were able to isolate and monitor plant physiological and microbial parameters under pre-industrial (200 ppm), ambient (400 ppm), and elevated (800 ppm) CO_2_ concentrations. The system enabled real-time quantification of net CO_2_ fluxes, revealing dynamic shifts in the balance between photosynthetic assimilation and plant and microbial respiration over time. Across all conditions, inoculation with the nodule symbiont *M. loti* significantly increased net CO_2_ uptake rates, particularly under elevated CO_2_, supporting the view that nitrogen availability becomes increasingly limiting as carbon input increases. This is in line with previous findings in *Medicago*, soybean and bean, where elevated CO_2_ enhances nodulation and the reliance on symbiotically fixed nitrogen *(13–15)*. Notably, this stronger CO_2_ drawdown is observed despite higher root-associated bacterial loads under elevated CO_2_ (**Fig. 4F**), which would be expected to increase belowground respiration and counteract net assimilation.Our results also raise the possibility that plant-microbial associations were important for the survival and performance of C3 plants during past low atmospheric CO_2_ periods, when carbon availability strongly constrained growth. Most low-CO_2_ studies report reduced growth, but rarely account for the root microbial community or for the ability of plants to shape that community. Our results, although still consistent with the idea that symbioses could be less important under low CO_2_, nonetheless suggest that microbial associations can remain a key determinant of plant performance even under carbon limitation.

Commensal bacteria alone had no detectable effect on plant growth but strongly influenced shoot-to-root carbon allocation, indicating that commensals alter host resource allocation even in the absence of canonical symbioses. When all three microbial groups (commensals, rhizobia, and AMF) were present, plants showed the most pronounced responses across nearly all traits, including biomass, nodulation, and CO_2_ flux. This pattern suggests that to fully benefit from elevated atmospheric CO_2_, plants may require multi-partner microbial associations that span trophic and functional niches. The importance of such synergistic interactions is underscored by the observation that the response of N-fixation to elevated CO_2_ is not universal, but can be positive, neutral, or even negative depending on the specific legume species and background soil nitrogen availability (*17*). Our work suggests that diverse microbial partnerships may help buffer against this variability, leading to a more robust plant response.

These findings are especially relevant given that many ecosystem-scale models that predict plant productivity and soil carbon cycling under future climate scenarios treat microbial activity as a black box, rarely accounting for the structure or composition of microbial communities (*19*). Our results point to a more complex interaction: not only do root-associated communities change in abundance and composition under elevated CO_2_, but the structure of microbial input (i.e., which microbial functional guilds are present) modulates plant responses at multiple levels and across multiple traits. Our measurements of root-derived carbon show that elevated CO_2_ increases carbon release from root systems and that the root microbiota determines the size of this response. In germ-free plants release rose 1.6-fold, with rhizobia and commensals present the increase was reduced, and abolished with commensals alone (**Fig. 4G**). Together with the increase in root bacterial loads under elevated CO_2_ (**Fig. 4F**), these patterns suggest that microbial partners determine the magnitude of carbon export under high CO_2_, retaining it in the root and associated microbes rather than releasing it, which in turn may amplify microbial activity and nutrient mobilization. Such feedback loops could significantly alter belowground carbon partitioning and nutrient fluxes, with consequences for carbon retention and release in soil ecosystems, since greater carbon input to an active microbiota under elevated CO_2_ accelerates the decomposition of soil organic matter and can limit soil carbon storage (*29, 30*).

Our data also highlight the unexpectedly strong influence of commensal microbes on plant growth traits. While commensals alone did not enhance growth, they dramatically influenced carbon allocation patterns, consistently driving higher root-to-shoot ratios, regardless of CO_2_ levels. This suggests that even in the absence of direct growth promotion, commensals can shift carbon allocation between root and shoot, possibly by preparing the root system for future symbiotic interactions or improving plant foraging capacity under nutrient-limited conditions. The underlying mechanism remains unclear but may involve the common symbiosis signaling pathway. This hypothesis is consistent with previous findings indicating altered commensal microbiota structures in symbiosis-impaired mutants in *L. japonicus* independently of nitrogen supply *(25, 31, 32)*. Whether this phenotype is driven by one or multiple members of the bacterial community remains unknown, but it appears to be a dominant and reproducible feature of commensal colonization.

Similarly, treatment with AMF showed significant interactions with other microbial partners. While AM fungi are often viewed as key players in plant nutrient acquisition, particularly phosphorus, their beneficial effects were negligible in our system, likely due to the high phosphate content of the sterile peat substrate used. However, when AMF were added alongside rhizobia and commensals, we observed further improvements across multiple traits, suggesting that their contribution may also depend on interactions with other microbiota members. This context-dependence of the beneficial impact of symbioses suggests that efforts to engineer plant microbiomes for future agriculture will require an understanding of microbial interkingdom interactions *(33)*, rather than relying on single-strain inoculants.

Beyond its specific findings, our study provides a scalable framework for studying plant-microbiota interactions under precisely controlled environmental conditions. The EcoChamber system enables quantitative integration of physiological, microbial, and gas exchange data, linking microbiota composition to plant carbon balance. Because the generation time of plant-associated microbes is orders of magnitude shorter than that of their hosts, the same system can be used to ask whether microbial adaptation to rising CO_2_ outpaces that of the plant over ecological timescales.

Taken together, our results demonstrate that the root-associated microbiota is an integral determinant of plant responses to altered CO_2_. In our experiments, the strongest responses across traits were observed when commensals, rhizobia, and AMF were all present under elevated CO_2_. Whether this pattern generalizes across plant species and soil types remains to be tested. Nonetheless, our findings imply that rising CO_2_ levels not only alter plant physiology, but also modify the structure and function of their microbial partnerships.

## Acknowledgments

This work was supported by the Max Planck Society and the Cluster of Excellence on Plant Sciences (CEPLAS), funded by the Deutsche Forschungsgemeinschaft (DFG, German Research Foundation) under Germany’s Excellence Strategy (EXC 2048/1, project no. 390686111). R.G.-O. acknowledges funding from the European Research Council (ERC) under the European Union’s Horizon Europe research and innovation programme (PHYCOSPHERES, grant agreement No. 101077231).

