## Supplementary material for "Symbiotic and commensal root microbes are key determinants of plant responses to rising CO_2_": Supllementary Information

**This PDF file includes:**

Materials and Methods

Supplementary Figures

Supplementary Tables

### Materials and Methods

#### *The EcoChamber system for gnotobiotic plant growth under varying $CO_2$ concentrations*

Growth experiments were performed in a controlled facility housing three identical climate cabinets (**Fig. 1B**; Hettich Benelux PRC1200 SL). All cabinets were operated under the same day/night regime (10 h light -14 h dark), light intensity (200 µmol $m^{-2}$ $s^{-1}$), temperature (21 °C), and air relative humidity (65%) for the duration of experiment. Environmental conditions inside each of these three cabinets differed only in their atmospheric $CO_2$ levels: 200 ppm, which received $CO_2$-depleted air via a $CO_2$ scrubber (CMC Instruments PG28L), 400 ppm, which remained at ambient atmospheric $CO_2$ concentration, and 800 ppm, which received $CO_2$-enriched air from gas cylinders (Air Liquide LASAL 2).

Each climate cabinet contained six separate EcoChamber units, custom-built using acrylic glass, with dimensions L x W x H = 322 x 167 x 212 mm. Specifications of the components used to build the system are listed in **Table S3**. The bottom and top shells were sealed from the outside environment by a circumferential, uninterrupted sealing gasket of silicone. Inside each EcoChamber, stainless steel racks (grade 1.4016) were placed to hold 20 FlowPots (*34*) filled with sterile peat growth substrate. The FlowPot system allowed the peat matrix to be inoculated with the desired microbial treatments (see Plant material and growth conditions section below) prior to sowing surface-sterilized, pre-germinated *L. japonicus* seedlings (three per individual FlowPot).

All EcoChambers received air through a hydrophobic filter (Sartorius Midisart 2000; 0.22 µm) at rate of ca. 2.5L/min. Upstream of this filter, a pneumatic valve-switching system and diaphragm ventilation pumps (Boxer 3112.660) were installed (**Fig. S4**). Within each EcoChamber, a perforated tube connected to the air inlet was installed, spanning the entire EcoChamber interior length. This element ensured that all FlowPots were equally exposed at to incoming air. Additionally, within each EcoChamber, a perforated tube connected to the air outlet was installed, spanning the entire EcoChamber interior length. This element ensured, that the outgoing air stream was exposed to a similar degree to all FlowPots. Air inflow was regulated by upstream solenoid valves of type 3/2-way (SMC VWD350) that directed the air either from the surrounding growth cabinet or through the sensor line back into the EcoChamber. Correspondingly, downstream solenoid valves of the same type diverted exhaust air from the EcoChamber either to the cabinet interior or to the sensor line, respectively. This valve arrangement enabled each EcoChamber to operate in either open-loop mode, when the air circulated between the EcoChamber and the cabinet, or closed-loop mode, when the air circulated between the EcoChamber and the sensor line **(Fig. 1** and **Fig. S4**).

At any given time during the experiments, a single EcoChamber was connected to the closed-loop circuit for 9 minutes while the remaining five operated in open-loop. The system cycled sequentially through all six EcoChambers in a 54-minute period. This design allowed all chambers to be measured every hour, while the 6 minutes left in the hour were used to measure the cabinet air through the sensor line. The sensor line (**Fig. 1** and **Fig. S4**) integrated a digital $CO_2$ sensor (Vaisala GMP343) using a single-beam dual-wavelength infrared principle, a thermal mass airflow sensor (Honeywell HAF), a piezoresistive absolute pressure sensor (WIKA A-10), a capacitive relative humidity sensor (TE Connectivity HM1500LF), and a platinum Pt1000-based temperature sensor (WIKA TF35) (**Fig. S8**). All components of the system are listed in **Supplementary Data 1**.

During the closed-loop mode, $CO_2$ concentrations inside each EcoChamber were measured through an infrared gas analyzer with an accuracy of ±3 ppm. Data from the sensor line were recorded every second. A proper utilization of the $CO_2$ sensor was achieved by considering three main functionality aspects: compensation, calibration, and configuration.

Compensation: the sensor obtains a value from its infrared detector, which is approximately proportional to the instantaneous quantity of $CO_2$ molecules within the sensor internal air cavity. In order to derive from it a value in ppm units, the manufacturer implemented a signal processing circuit which takes into account values of pressure, temperature, humidity and oxygen content in a process defined as compensation. The result of this process is a relative quantity in ppm, describing the fraction of $CO_2$ molecules to all molecules within the sensor cavity. Of the four above-mentioned compensation values, the temperature was continuously updated by a dedicated built-in temperature sensor, while for pressure, humidity and oxygen, constant default values were used (1013.250 hPa, 50%, 20.95% respectively). When interfacing with the sensor, the user thus received a stream of $CO_2$ values already translated in ppm units.

Calibration: despite the benefits of the dual-wavelength operating principle, the accuracy of the $CO_2$ sensor deteriorates over time. The sensor thus offers a calibration mode over a serial bus interface, during which it is temporarily exposed to an environment of a pre-determined $CO_2$

concentration. This is achieved by gas cylinders containing reference gas, at $CO_2$ levels of e.g. 0ppm, 500ppm or 1000ppm. In order to avoid disrupting the system between the experiments, it was decided to conduct a calibration only at the termination of the experimental campaign. At that instance, the maximum observed relative error between sensor reading and calibration gas was ca. 8%. Since the flux calculation is performed by using the difference between values of the same sensor unit, it was decided that the propagation of the absolute error to the flux calculation was low. The $CO_2$ levels inside the climate cabinet were regulated by a different set of sensors.

Configuration: the user can tailor the temporal characteristics of the sensor value stream over a serial bus interface, by individually setting the acquisition period, the strength of the median filter averaging filter and smoothing filter. For the experiment, the values of 1s, 10 points, 30 points and 0 points (disabled) were used, respectively.

At the end of the closed-loop mode $CO_2$ concentration, temperature, and pressure data were used to calculate the rate of $CO_2$ assimilation or release. Raw $CO_2$ data were expressed as the ratio of $CO_2$ molecules to the total number of air molecules in the circulating volume. To convert these relative values into absolute molar quantities, the concentration in ppm was transformed using the ideal gas law:

$$n_{CO_2}\text{ (mol)} = CO_2\text{ (ppm)} \times n_{\text{total}}\text{ (mol)} = CO_2\text{ (ppm)} \times (p \times V) / (R \times T)$$

where $CO_2$ is the $CO_2$ concentration (ppm, obtained by the $CO_2$ sensor), p is the air pressure (Pa, obtained by the pressure sensor), V is the volume of the circulating air ($m^3$, obtained by the geometry of the circulating air volume), R is the ideal gas constant (J $molK^{-1}$), and T is the air temperature (K, obtained by the temperature sensor).

The resulting $CO_2$ quantity (µmol) was expressed as a time series, and the rate of change (µmol $h^{-1}$) was estimated using the Theil-Sen slope estimator. The following equation describes the Theil-Sen algorithm for the estimation of the first derivative of $CO_2$ concentration with respect to time:

$$S_{ij} = (\Delta n / \Delta t)_{ij} = (n_j - n_i) / (t_j - t_i)$$

where $S$ is the matrix of pairwise slopes (µmol $s^{-1}$) for a pair of time points with indices $i$ and $j$, each in the range of 1-420 (corresponding to the 7-minute measurement window that follows a 2-minute flushing period at the start of the closed-loop mode), $n$ is the concentration of $CO_2$ (µmol) at the time point specified by $i$ or $j$, and $t$ is the time specified by $i$ or $j$ (s). For each measurement series, we calculated the corresponding pairwise slopes, converted them to a sorted vector, and selected the median value as the robust first derivative of $CO_2$ concentration with respect to time, representing the net $CO_2$ uptake or release rate.

EcoChambers were also equipped with an automatic irrigation system, comprising a 20-fold multi-port internal manifold (Formlabs BioMed Resin) attached to silicone tubes arranged to ensure that each pot would receive approximately the same amount of sterilized $ddH_2O$ (**Fig. 1**). All components of the irrigation system were sterilized by autoclaving, UV light, or ethanol depending on the material specification. Sterilized $ddH_2O$ was delivered to the EcoChambers through a geared water pump (TSC MGD1000S) connected to a water bottle and to 6 irrigation valves of type 2/2-way (SMC VDW22LA) through a common external manifold. Between the valves and the EcoChamber a non-return valve (Bürkle 229-0134) and a hydrophilic filter (Sartorius Sartolab P20, 0.22 µm) were placed to eliminate the possibility of flow-back and cross-contamination (**Fig. S5**).

The EcoChamber system was designed to integrate sensing, actuation, and telemetry within a single automated framework for environmental monitoring and control (**Fig. S7**). Sensor data and actuator control signals were managed through a structured electronic and software architecture to ensure precise and synchronized operation. The system layout included dedicated electrical and communication pathways for signal flow and power distribution. Sensor signals were transmitted through designated circuits, while power distribution was organized by positive or negative voltage load, ground line, and protective earth connections.

Sensor values and LED status information from the climate cabinet system were transmitted every five seconds through the Ethernet interface (ETH0) to a central Linux server (version 5.10.0-18-amd64 #1 SMP Debian 5.10.140-1). In parallel, sensor data from the embedded EcoChamber system were transmitted every second through a separate Ethernet interface (ETH1) by the microcontroller board (µCU0; STMicroelectronics Nucleo-H723ZG) This microcontroller was equipped with a base shield (Grove Base Shield V2 Arduino), multiple analog and digital interface ports, enabling sensor acquisition, actuator control and telemetry handling. It managed both the automatic switching of the solenoid air valves and the digital signal processing of sensor inputs. The embedded software controlling these operations was implemented in C using the Zephyr real-time operating system (version 3.2.0). Verification of the microcontroller functionality was conducted with the software tool Minicom (version 2.9), which enabled status and error messages to be read from a computer over USB protocol. The raw data were collected from the server by means of a Rest client (version 3.6.2), without interfering with the data recording process.

Telemetry data from all connected components were routed via a common Ethernet switch to the Linux server, where a Docker (version 20.10.5) container running ThingsBoard (version 3.6.4) enabled continuous remote monitoring and visualization of system parameters. The EcoChamber was powered by a single 230 V alternating current (AC) source, which fed multiple direct voltage (DC) converters providing 5 V, 12 V, and 24 V outputs to meet the voltage requirements of individual components (**Fig. S6** and **Fig. S7**). Airflow regulation for each EcoChamber was achieved through potentiometers (Bourns 3296W-1-102), powered by a voltage reference (STMicroelectronics L7805A) that controlled airflow for two chambers simultaneously (**Fig. S8**). An additional potentiometer was used to regulate airflow for ambient air measurements, ensuring consistent environmental monitoring conditions across all experimental units. The voltage reference unit was equipped with an aluminum fin for transferring the dissipated heat to the ambient air.

***Bacterial and AMF material***

Bacterial strains composing the Synthetic Community (SynCom) in this study were acquired from the *Lj* bacterial culture collection (*25*) and are listed in **Table S1**. The strains were grown in tryptic soy broth (15 g $l^{-1}$, TSB, Sigma-Aldrich) on agar plates containing 15 g $l^{-1}$ of Bacto Agar (Difco) for 5 days followed by incubation in TSB liquid medium for 72 h at 25°C and 180 rpm.

The *M. loti* strain (LjRoot218) was cultivated in TY medium (5 g $l^{-1}$ tryptone, 3 g $l^{-1}$ yeast extract, 0.5 g $l^{-1}$ $K_2HPO_4 \cdot 3H_2O$, 0.2 g $l^{-1}$ $MgSO_4 \cdot 7H_2O$, 0.1 g $l^{-1}$ NaCl), on agar plates and in liquid medium.

*Rhizophagus irregularis* DAOM 197198 AMF spores (Agronutrition) were washed in LAS medium (0.294 g $l^{-1}$ $CaCl_2$, 0.085 g $l^{-1}$ $NaNO_3$, 0.174 g $l^{-1}$ $K_2SO_4$, 0.184875 g $l^{-1}$ $MgSO_4 \cdot 7H_2O$, 0.0005696 g $l^{-1}$ $Na_2HPO_4 \cdot 2H_2O$, 0.009175 g $l^{-1}$ FeNa-EDTA, 0.000845 g $l^{-1}$ $MnSO_4 \cdot H_2O$, 0.000062375 g $l^{-1}$ $CuSO_4 \cdot 5H_2O$, 0.00014375 g $l^{-1}$ $ZnSO_4 \cdot 7H_2O$, 0.00155 g $l^{-1}$ $H_3BO_3$, 0.0000242 g $l^{-1}$ $Na_2MoO_4 \cdot 2H_2O$, 0.0000562 g $l^{-1}$ $CoSO_4 \cdot 7H_2O$) to remove the non-germination buffer and incubated in the same medium for 72 h with $10^{-3}$ µM rac-GR24 (Chiralix) to induce germination.

After the incubation period, bacterial strains were washed twice in 10 mM $MgSO_4$, the maximum *in vitro* growth rate of each strain was determined using a plate reader (Tecan Infinite 200 PRO) to adjust the cultures to an initial $OD_{600}$ of 0.02 and the strains were mixed according to desired synthetic community (SynCom). Germinated AMF spores were also washed twice in 10 mM $MgSO_4$, their concentration was verified in a dilution of 1:100, and finally the spores were mixed with bacterial strains in LAS medium or diluted in LAS medium alone.

***Plant material and growth conditions***

*L. japonicus* accession Gifu B-129 was used as plant host in this study. *Lotus* seeds were scarified with sandpaper, incubated in 1% bleach for 20 minutes, rinsed eight times with sterile $ddH_2O$, and incubated in $ddH_2O$ for 2 hours. After this period, the sterilized seeds were rinsed once more and germinated on sterile Whatman paper moistened with $ddH_2O$ in square Petri dishes for 72 h (10 h light at 21 °C and 14 h dark at 21 °C).

All experiments were conducted inside the EcoChambers that had been exposed to UV light overnight. Plants were grown in a modified FlowPot system (*34*; peat:vermiculite ratio, medium and housing differ from the published protocol), which was placed inside the EcoChamber after sowing the germ-free *L. japonicus* seedlings. The peat substrate used in this experiment was mixed with vermiculite in a 2:1 ratio, moistened with 200 ml $ddH_2O$ per liter, and autoclaved once daily for 3 days. FlowPots were assembled by filling each pot with 65 ml of autoclaved soil mixture and a nylon mesh placed in the bottom to prevent clogging of the system and autoclaved again once daily for another 3 days. After the last autoclaving, FlowPots were flushed with 50 ml of sterile $ddH_2O$ via vacuum infiltration using a QIAvac 24 Plus manifold (Qiagen). Next, FlowPots were flushed with 50 ml of sterile LAS medium (axenic condition) or LAS medium supplemented with the microbial inoculum.

The FlowPots were then inserted in their corresponding EcoChambers, where the plants were cultivated for 5 weeks. All procedures mentioned above were conducted under the clean bench to avoid any contamination. Sterility of the autoclaved substrate was verified before each experiment by plating samples of the soil mixture, on which no growth was observed. Each EcoChamber housed 20 FlowPots with a single microbial treatment or the axenic condition and was placed in the plant growth cabinet corresponding to the desired $CO_2$ concentration. Once per week, 200 ml of sterilized $ddH_2O$ was provided to the EcoChambers. This volume was determined by extensive preliminary tests of evaporation rate and water retention of the soil matrix. The beforementioned air humidity control of the climate cabinet which housed the

EcoChambers ensured that the soil did not dry between irrigation cycles. After the 5-weeks growth period, plants were destructively harvested and the following traits were measured: shoot fresh weight, shoot length, chlorophyll content, root fresh weight, root-to-shoot ratio, number of nodules, and total weight of nodules. Nodules were separated from roots with a scalpel before weighing, so root fresh weight and the root-to-shoot ratio exclude nodules. Nodule number and total nodule weight were recorded separately. Roots, leaves, and rhizosphere material were collected and immediately flash-frozen in liquid nitrogen for subsequent DNA extraction using the ZymoBIOMICS DNA Miniprep Kit (D4300).

#### *16S rRNA amplicon sequencing*

For each microbial treatment containing bacterial strains and for all $CO_2$ concentrations, bacterial community composition was assessed by amplicon sequencing of the 16S rRNA gene. The 16S rRNA V5-V7 region was amplified using primers 799F and 1192R under the following conditions: 94 °C for 2 min; 25 cycles of 94 °C for 30 s, 55 °C for 30 s, and 72 °C for 30 s; followed by a final extension at 72 °C for 10 min (*33*). PCR products were treated with 1 µl Antarctic phosphatase, 1 µl Exonuclease I, and 2.44 µl Antarctic Phosphatase buffer (New England BioLabs GmbH, Frankfurt, Germany). A second PCR was performed to attach sample-specific barcodes using 3 µl of the digested PCR product as template, with the same cycling conditions, except for 10 cycles. Amplicon size and quality were verified by gel electrophoresis. Following the second PCR, DNA concentrations were measured, and all samples were pooled to generate the sequencing library. Gel electrophoresis and QIAquick gel extraction (QIAGEN) were used to remove chloroplast-derived 16S rRNA fragments. The final library was purified with AMPure XP beads (Beckman Coulter) and eluted in 30 µl of nuclease-free water. DNA concentrations were quantified using the QuantiFluor system (Promega), and sequencing was carried out on the Illumina MiSeq platform.

#### *Processing of 16S rRNA gene amplicon data*

Raw sequencing reads obtained from the MiSeq platform were demultiplexed using QIIME2 v2021.11.0 (*35*). Forward and reverse reads were subsequently merged with FLASH v2.2.00 (*36*). Sequences containing undetermined nucleotides (i.e., “N” bases) were identified with usearch v10.0.240 (*37*) and removed from the dataset. Reference-based error correction of high-quality sequences was performed using Rbec v1.1.4 (*38*) and a phred-scaled quality threshold of 33. The resulting count table was corrected for 16S rRNA gene copy-number variation and filtered for a minimum depth of 1,000 reads. Subsequent analyses were conducted in R. Counts were normalized by calculating strain-specific relative abundances after *in silico* removal of *M. loti*. PCoA (**Fig. 4A**) and CPCoA (**Fig. 4B-C**) were computed with the capscale function of the vegan package, and PCoA and CPCoA plots, barplots (**Fig. 4D**), and line plots (**Fig. S2A-B**) were generated using ggplot2 v4.0.3.

#### *Quantification of bacterial load and AMF colonization on plant roots*

Bacterial load on plant roots was quantified by determining the relative abundance of bacterial to plant DNA using quantitative PCR (qPCR). For bacterial detection, the V5-V7 region of the 16S rRNA gene was amplified with primers 799F (AACMGGATTAGATACCCKG) and 1192R (ACGTCATCCCCACCTTCC). For *L. japonicus* Gifu B-129, a fragment of the *Nfr5* gene was amplified using primers TCATATGATGGAGGAGTTGTCTGTT and ATATGAGCTTCGGAGCATGG. qPCR was performed as previously described (*25*). The bacterial abundance in each sample was expressed as the ratio of 16S rRNA gene copies to *Nfr5* gene copies, calculated according to the equation:

$$(16S\ rRNA) / (\mathit{Nfr5}) = 2^{-Ct(16S)} / 2^{-Ct(\mathit{Nfr5})}$$

For AMF detection, a region of the mitochondrial DNA was amplified with primers 197198F (CCCACCAGGGCAGATTAATC) and 197198R (TGGCTTTGTACAGGCAACAG) (*39*) and further analyzed in the same way as the 16S rRNA gene.

***Exudates collection and TOC measurements***

Plants were grown as described in the section above (Plant material and growth conditions). For this experiment axenic, SC and SC + *M. loti* treatments were used to test whether total organic carbon (TOC) released by *L. japonicus* roots changes between ambient (400 ppm) and elevated (800 ppm) $CO_2$ and whether this change depends on microbial associations. The experiment was performed twice, with axenic and SC + *M. loti* plants in both runs and SC plants in the second run only. On day 34 of plant growth, EcoChambers were removed from the growth cabinets and transferred to a clean bench. Roots were carefully harvested, thoroughly washed to remove peat matrix particles, and transferred to new FlowPots containing 35 mL of autoclaved dd$H_2O$. This volume was selected based on preliminary tests using different volumes to maximize root-water contact. FlowPots were sealed at the bottom with a LuerLock connector (Cole-Parmer 45501-28) and fitted with a 3D-printed collar (Formlabs RS-F2-BMBL-01) at the top to secure the shoot, ensuring that only roots were in contact with the solution. The assembled FlowPots were placed in gnotobiotic boxes (Sac $O_2$ TP5000 and TPD5000) and returned to their respective $CO_2$ growth cabinets. After 24 h, root exudates were collected into Falcon tubes by opening the LuerLock at the base of the FlowPot and stored at -20 °C until further analysis. TOC in the collected exudates was quantified using a DIMATOC 2000 analyzer (Dimatec) as the difference between total and inorganic carbon. The amount of carbon released per pot was calculated as the TOC concentration multiplied by the collected volume. Because absolute amounts differed between experiments, values were normalized to the median of the 400 ppm pots of the same treatment within each experiment before the two experiments were combined. Differences between $CO_2$ levels within each treatment and between treatments at 800 ppm were tested with Wilcoxon rank-sum tests. Fold changes are ratios of medians with bootstrap 95% confidence intervals.

***Statistics and reproducibility***

All experiments were performed with full factorial (biological and technical) replication. The complete design was repeated in four independent experiments (A to D in the figures), each run with freshly prepared inocula and substrate, so that every comparison between treatments or $CO_2$ levels is replicated across experiments. Whenever bacterial abundances or plant growth parameters were compared, we first assessed the data distribution. In the case of a normal distribution, an ANOVA test was performed, followed by Tukey's test. In the case of non-normal distribution, a non-parametric Kruskal-Wallis test was performed, and whenever appropriate, *P* values were adjusted for multiple testing using the Benjamini-Hochberg method ($\alpha = 0.05$).

Statistical tests on beta-diversity analyses were performed using a PERMANOVA test (vegan) with 5,000 random permutations. Whenever boxplots were used in figures, data were represented as median values (horizontal line), Q1 - 1.5× interquartile range (boxes) and Q3 + 1.5× interquartile range (whiskers).

#### *Code and data availability*

Raw sequencing reads have been deposited at the European Nucleotide Archive (ENA) under accession numbers PRJEB102711 (amplicon sequencing). Scripts for raw data processing, analysis, and plotting are available on GitLab: https://gitlab.gwdg.de/l.anastasopoulos/lotus-data and https://github.com/marina-cotta/Symbiotic-and-commensal-root-microbes-are-key-determinants-of-plant-responses-to-rising-CO-.

**Supplementary Figures**

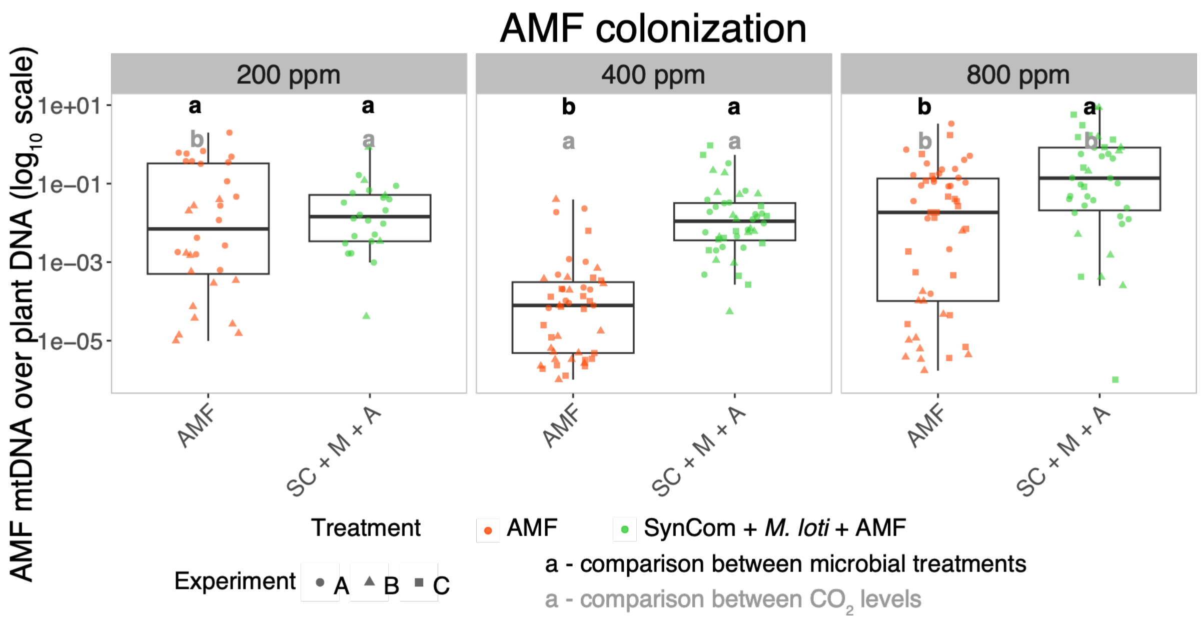


**Figure S1**. **AMF abundance in roots systems**. AMF colonization in root systems quantified by qPCR of a mitochondrial genome region of *R. irregularis* DAOM-197198 normalized to the L. japonicus *Nfr5* gene. The y axis is on $log_{10}$ scale. Each data point reflects qPCR results from 3 pooled plants per replicate (*n* = 60 plants per treatment group). Statistical significance was determined by Kruskal-Wallis or ANOVA

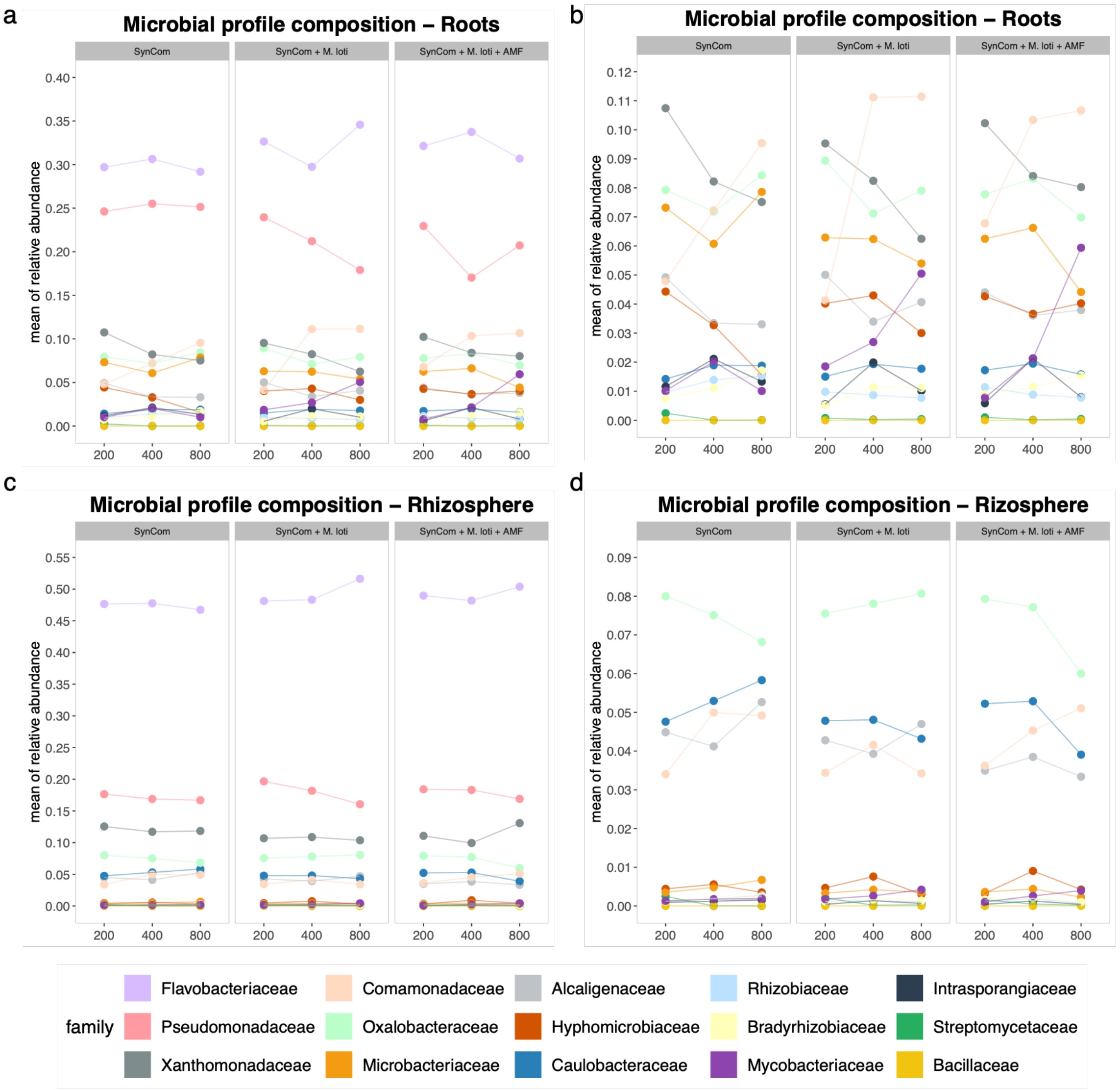


**Figure S2**. Compositional profiles of SynCom retrieved from roots and rhizosphere under increasing CO2 levels. Microbial profile of SynCom members from roots (**A**) and rhizosphere (**B**) for each microbial treatment under different $CO_2$ conditions. Secondary plots with adjusted scales highlight changes in the relative abundance of low-abundant SynCom members in roots and rhizosphere across CO2 levels.

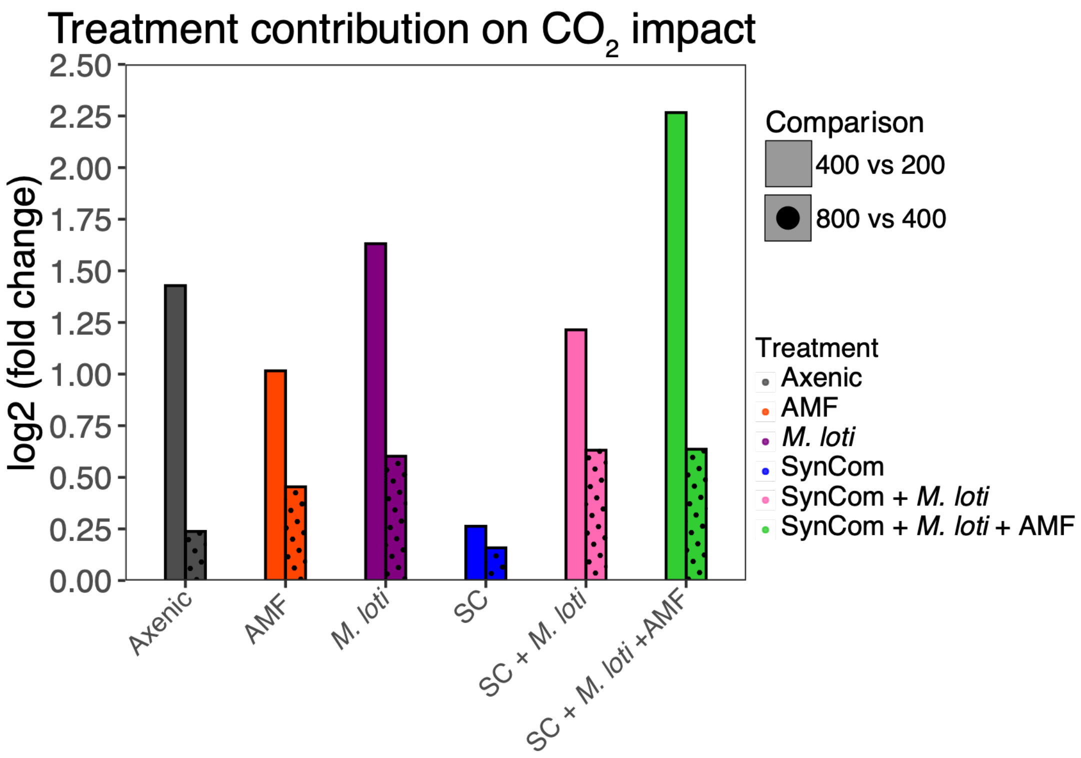


**Figure S3**. **Contributions of microbial treatment on the impact of rising $CO_2$ levels on *L. japonicus*.** Bar plots showing the $CO_2$-dependent response of *L. japonicus* to different microbial treatments based on ten traits: shoot fresh weight, shoot length, chlorophyll content, root fresh weight, root-to-shoot ratio, nodule number, total nodule weight, bacterial abundance, and $CO_2$ fluxes. Each bar represents the average fold change for these traits, comparing $CO_2$ levels of 200 to 400 ppm (dotted bars) and 800 to 400 ppm (solid bars) for a specific microbial treatment.

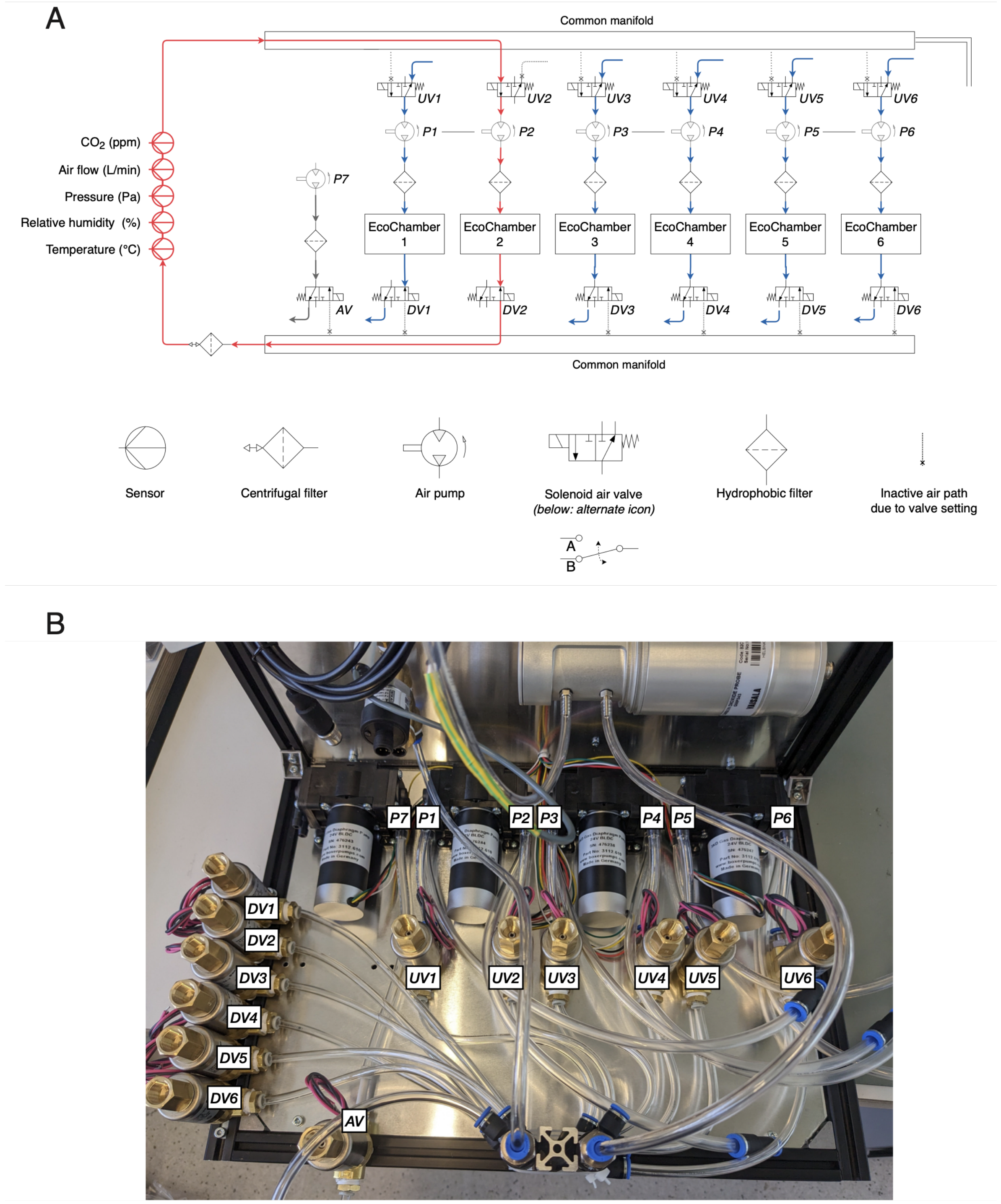


**Figure S4**. **Process diagram of EcoChamber ventilation.** Process diagram of the complete ventilation system for EcoChamber air supply. The diagram depicts exemplary EcoChamber EC2 in closed-loop mode and utilizing the sensor line, while the remaining five EcoChambers are in open-loop mode. The air pumps for the corresponding to the respective EcoChambers are labeled as P1-P6. The solenoid air valves upstream of the EcoChambers are labeled as UV1-UV6, whereas the solenoid air valves downstream of the EcoChambers are labeled as DV1-DV6. The air pump and solenoid air valve for the ambient measurement is labeled as P7 and AV respectively. Each EcoChamber receives air through a hydrophobic filter. The sensor line is protected by a common centrifugal filter.

A

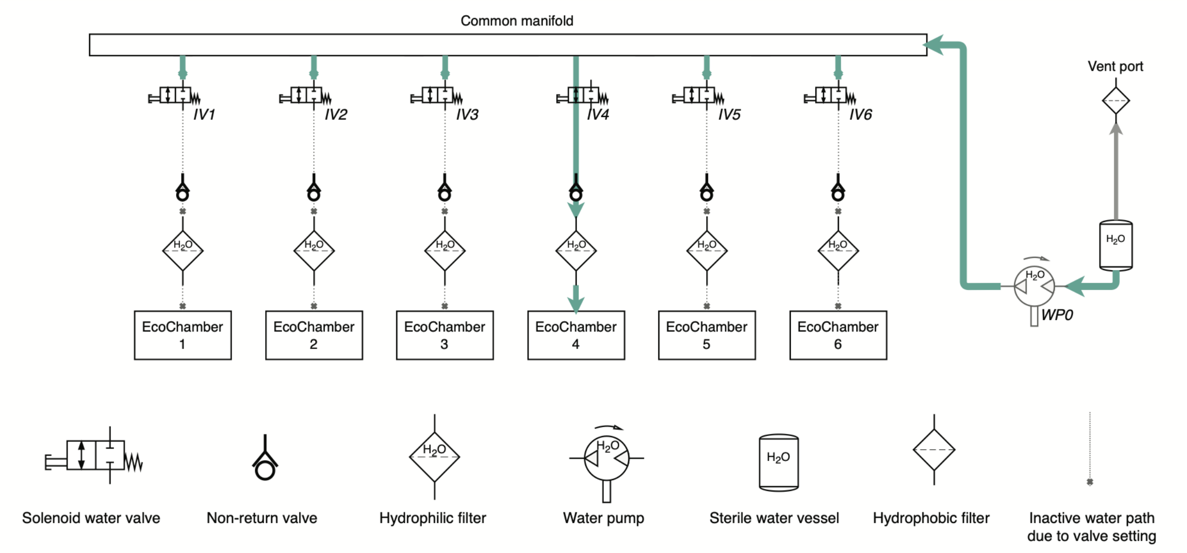


B

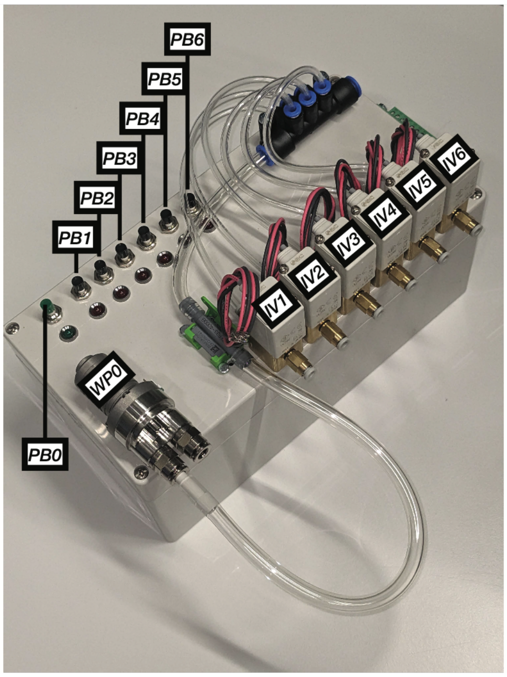


**Figure S5**. **Process diagram of EcoChamber irrigation**. Process diagram of the complete irrigation system for EcoChamber water supply. The top and bottom schematics separately depict the flow of water and electricity respectively during irrigation, for the same system. Dark green color indicates the flow of water, whereas red and black color indicate the flow of electricity flow. In the latter case red is used for the positive side of the load voltage, whereas black is used for the negative/ground side of the load voltage. Additionally, light green color indicates the protective earth wiring. The water irrigation pump (WP0) is physically connected to the six irrigation valves (IV1 to IV6) by a common manifold. The diagram exemplary depicts EcoChamber EC4 during irrigation. Push button PB0 is used to engage the water irrigation pump with a direct voltage of 24V, whereas push buttons PB1 to PB6 are used to engage the respective irrigation valves with a direct voltage of 5V. A common alternate voltage source of 230V supplies electricity to the two mentioned low voltage sources.

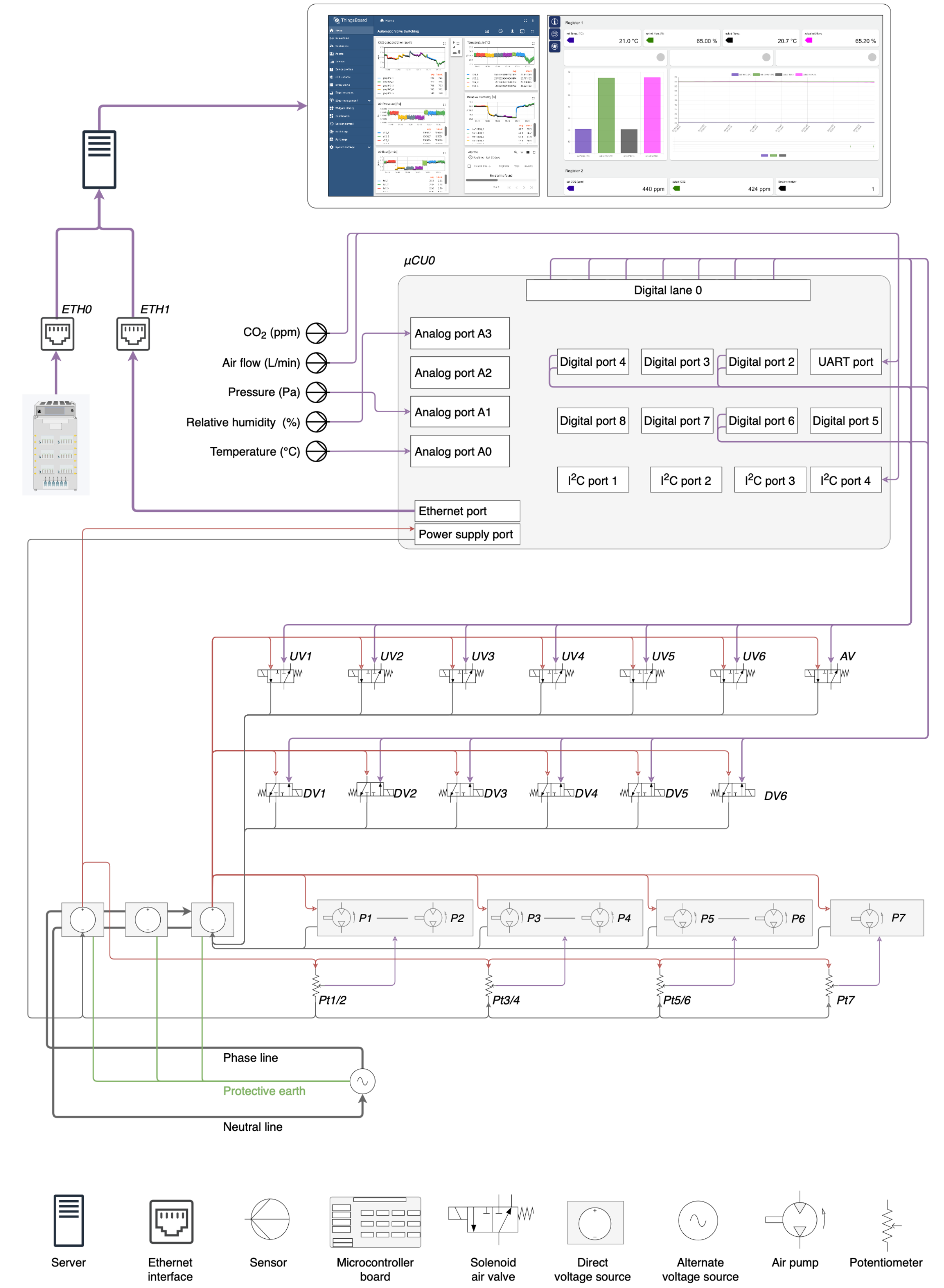


**Figure S6**. **Process diagram of EcoChamber sensing, actuation and telemetry.** Process diagram of the complete EcoChamber system for sensing, actuation and telemetry. Purple color is used for the flow of sensor signals, whereas red and black color are used for the electricity flow. In the latter case red is used for the positive side of the load voltage, whereas black is used for the negative/ground side of the load voltage. Additionally, light green color indicates the protective earth wiring. Once every 5 seconds, sensor values and LED status information from the climate cabinet system are published through Ethernet interface ETH0 to a Linux server. Once every 1 second, sensor values from the embedded EcoChamber system are published through Ethernet interface ETH1 to the same Linux server by Microcontroller board µCU0. This microcontroller also manages the automatic switching of the solenoid air valves and the digital signal processing of the sensor signals. The software is written in C++ language utilizing the Zephyr real-time operating system. Onto the Linux Server (which receives the telemetry data packages from a common Ethernet switch), a Docker container with Thingsboard functionality is installed, enabling remote system monitoring. A single 230V alternate voltage source provides the electrical power to the EcoChamber system. Multiple direct voltage sources (for 5V, 12V, 24V) direct the electricity to specific target devices, depending on the respective voltage requirements. Potentiometers Pt1-Pt6 are used for regulating the air flow for the respective EcoChambers. The regulation is conducted in pairs of two, since one potentiometer serves two pump chambers at the same time. Potentiometer Pt7 is used for regulating the air flow for the ambient measurement.

### Supplementary Tables

**Table S1. Composition of the bacterial SynCom used in this study.** Strains were selected from the *Lj*-SPHERE culture collection based on taxonomic coverage and the relative abundance of their taxonomic groups in natural communities (Wippel *et al.*, 2021).

| Strain ID | Phylum | Class | Order | Family | Genus |
|---|---|---|---|---|---|
| LjRoot1 | Proteobacteria | Betaproteobacteria | Burkholderiales | Alcaligenaceae | *Achromobacter* |
| LjRoot5 | Firmicutes | Bacilli | Bacillales | Bacillaceae | *Bacillus* |
| LjRoot222 | Proteobacteria | Alphaproteobacteria | Rhizobiales | Hyphomicrobiaceae | *Devosia* |
| LjRoot17 | Proteobacteria | Alphaproteobacteria | Caulobacterales | Caulobacteraceae | *Brevundimonas* |
| LjRoot24 | Actinobacteria | Actinobacteria | Actinomycetales | Intrasporangiaceae | n.d. |
| LjRoot33 | Proteobacteria | Betaproteobacteria | Burkholderiales | Oxalobacteraceae | *Herbaspirillum* |
| LjRoot44 | Actinobacteria | Actinobacteria | Actinomycetales | Microbacteriaceae | *Microbacterium* |
| LjRoot46 | Proteobacteria | Alphaproteobacteria | Rhizobiales | Rhizobiaceae | *Shinella* |
| LjRoot52 | Proteobacteria | Alphaproteobacteria | Rhizobiales | Bradyrhizobiaceae | *Bosea* |
| LjRoot59 | Proteobacteria | Gammaproteobacteria | Pseudomonadales | Pseudomonadaceae | *Pseudomonas* |
| LjRoot60 | Proteobacteria | Gammaproteobacteria | Xanthomonadales | Xanthomonadaceae | *Pseudoxanthomonas* |
| LjRoot72 | Proteobacteria | Betaproteobacteria | Burkholderiales | Comamonadaceae | *Acidovorax* |
| LjRoot80 | Actinobacteria | Actinobacteria | Actinomycetales | Mycobacteriaceae | *Mycobacterium* |
| LjRoot82 | Bacteroidetes | Flavobacteriia | Flavobacteriales | Flavobacteriaceae | *Flavobacterium* |
| LjNodule218 | Proteobacteria | Alphaproteobacteria | Rhizobiales | Phyllobacteriaceae | *Mesorhizobium* |
| LjRoot303 | Actinobacteria | Actinobacteria | Actinomycetales | Streptomycetaceae | *Streptomyces* |

**Table S2. Soil nutrient composition.** Nutrient levels in the peat matrix were measured before seedling sowing and after cultivation under the different microbial treatments and $CO_2$ concentrations. Values are reported in ppm for all components, except nitrate, which is expressed as a percentage.

| Treatment | $CO_2$ (ppm) | pH | Ec (mS/cm) | Organic matter (%) | P (ppm) | K (ppm) | Mg (ppm) | Ca (ppm) | Na (ppm) | Fe (ppm) | Zn (ppm) | Mn (ppm) | Cu (ppm) | B (ppm) | Total Nitrogen (%) |
|---|---|---|---|---|---|---|---|---|---|---|---|---|---|---|---|
| Axenic | 200 | 5.84 | 1.988 | 26.5 | 25.02 | 560.5 | 1761.66 | 8899.54 | 224.55 | 54.6 | 6.12 | 57.79 | 16.45 | 9.39 | 0.307 |
| Axenic | 400 | 5.51 | 1.816 | 16.5 | 12.27 | 376.71 | 1889.84 | 8168.75 | 154.24 | 39.92 | 3.33 | 35.6 | 11.51 | 8.22 | 0.254 |
| Axenic | 800 | 6.22 | 1.442 | 24 | 19.3 | 591.3 | 2062.04 | 11665.9 | 284.15 | 62.48 | 4.39 | 47.11 | 13.14 | 6.31 | 0.287 |
| AMF | 200 | 6.27 | 1.362 | 25.5 | 19.43 | 464.31 | 2154.49 | 11047.4 | 165.76 | 39.43 | 4.63 | 40.62 | 13.07 | 7.5 | 0.234 |
| AMF | 400 | 5.36 | 2.002 | 20 | 11.18 | 333.02 | 1609.48 | 8150.38 | 127.19 | 38.92 | 3.5 | 35.48 | 10.28 | 8.37 | 0.243 |
| AMF | 800 | 5.95 | 1.739 | 18.5 | 18.3 | 534.53 | 1834.7 | 9897.48 | 274.01 | 54.55 | 6.06 | 45.25 | 11.81 | 7.98 | 0.32 |
| *M. loti* | 200 | 5.85 | 1.851 | 28 | 20.67 | 538.54 | 2128.26 | 10250.6 | 207.09 | 44.5 | 5.05 | 47 | 15.32 | 14.49 | 0.287 |
| *M. loti* | 400 | 5.52 | 1.755 | 18 | 8.85 | 400.47 | 2366.28 | 9406.78 | 268.05 | 36.14 | 2.95 | 30.52 | 11.19 | 6.83 | 0.296 |
| *M. loti* | 800 | 5.64 | 1.231 | 15 | 13.97 | 235.81 | 1286 | 6641.21 | 107.78 | 57.33 | 3.97 | 35.59 | 10.07 | 6.3 | 0.212 |
| SynCom | 200 | 5.93 | 1.81 | 25.5 | 25.59 | 610.78 | 2265.94 | 11176.2 | 188.75 | 42.01 | 4.88 | 44.03 | 13.8 | 10.75 | 0.252 |
| SynCom | 400 | 5.46 | 1.776 | 17 | 13.43 | 431.71 | 2110.53 | 8453.22 | 238.67 | 48.89 | 3.72 | 41.95 | 13.7 | 7.94 | 0.252 |
| SynCom | 800 | 6.12 | 1.442 | 27.5 | 18.92 | 720.24 | 2179.84 | 12053.6 | 540.63 | 55 | 3.95 | 43.16 | 11.51 | 8.07 | 0.235 |
| SynCom + *M. loti* | 200 | 5.88 | 1.735 | 25.5 | 24.17 | 539.11 | 2220.28 | 11405.2 | 176.28 | 41.24 | 5.17 | 40.46 | 12.42 | 9.99 | 0.27 |
| SynCom + *M. loti* | 400 | 5.54 | 1.708 | 18.5 | 11.03 | 317.44 | 1903.61 | 7627.19 | 200 | 41.98 | 3.33 | 35.79 | 12.95 | 7.37 | 0.223 |
| SynCom + *M. loti* | 800 | 5.81 | 1.284 | 16.5 | 14.51 | 429.74 | 2132.88 | 10475.2 | 139.62 | 57.01 | 3.56 | 37.25 | 11.69 | 7.51 | 0.18 |
| SynCom + *M. loti* + AMF | 200 | 6.13 | 1.435 | 20 | 23.44 | 579.49 | 2354.15 | 12053.3 | 212.06 | 43.47 | 7.77 | 43.77 | 12.77 | 6.86 | 0.34 |
| SynCom + *M. loti* + AMF | 400 | 5.38 | 1.708 | 18.5 | 11.9 | 328.94 | 2129.45 | 9054.8 | 357.81 | 45.29 | 3.8 | 38.41 | 11.64 | 7.63 | 0.287 |
| SynCom + *M. loti* + AMF | 800 | 5.83 | 1.71 | 18.5 | 21.39 | 420.1 | 1742.69 | 9652.71 | 161.88 | 60.46 | 4.77 | 43.01 | 12.3 | 8.59 | 0.276 |
| Soil before plant growth | NA | 5.91 | 1.412 | 21 | 25 | 693.08 | 1784.69 | 8714.16 | 170.05 | 70.52 | 4.68 | 51.58 | 11.83 | 10.28 | 0.311 |

**Table S3. EcoChamber material table.** A table containing all items used for building and commissioning the EcoChamber system, sorted in categories. The material list contains the manufacturer or supplier name, article number, technical specifications and reference to the figures and schematics of the manuscript depicting the respective item

| Category | Description | Manufacturer | Article number | Specification | Figure reference | Label reference | Microelectronics reference |
|---|---|---|---|---|---|---|---|
| Enclosure | Climate cabinet | Hettich Benelux | PRC1200 SL | Volume: 968 L | 1 | n/a | n/a |
| Enclosure | EcoChamber | GöLi Glas & Kunststoffe | n.a. (custom design) | Material: acrylic Plexiglass, Volume: 10 L | 1 | n/a | n/a |
| Enclosure | EcoChamber gasket | Plazura | 2023-17291 | Material: silicone foam | 1 | n/a | n/a |
| Enclosure | EcoChamber inlet port | Landefeld | KQ2E06-00A | Material: Brass | 1 | n/a | n/a |
| Enclosure | EcoChamber toggle latches | Otto Ganter | GN 832.2-55-A-ST | Material: Steel, Zinc plated, blue passivated | 1 | n/a | n/a |
| Enclosure | EcoChamber flow pot tray | Wilms Metallmarkt | n.a. (custom design) | Material: Stainless steel 1.4016 | 1 | n/a | n/a |
| Electrical supply | Enclosure for electronic equipment | ABB | Mistral 65, 1SLM006500A1213 | Protection class: IP65 | n/a | n/a | n/a |
| Electrical supply | Enclosure power provision | RS | 190-4190 | Output voltage: 5 V, Output power: 15 W | S7 | n/a | n/a |
| Electrical supply | Enclosure power provision | RS | 190-4191 | Output voltage: 12 V, Output power: 24 W | S7 | n/a | n/a |
| Electrical supply | Enclosure power provision | RS | 190-4193 | Output voltage: 24 V, Output power: 36 W | S7 | n/a | n/a |
| Control electronics | Microcontroller unit | STMicroelectronics | Nucleo H723ZG | STM32H723ZG MCU | S7 | µCU0 | n/a |
| Control electronics | Base shield | Seed Technology | 103030000 | Type: Grove Base Shield V2 Arduino | n/a | n/a | n/a |
| Sensing | Pin header | Degson | DG308-2.54-02P-14-00AH-1 | Grid: 2.54mm | n/a | n/a | n/a |
| Sensing | Air CO2 concentration - sensor module | Vaisala | GMP343B2C1A0N1N0A | Range: 0…2000 ppm | 1, S4, S8 | n/a | J2 |
| Sensing | Air CO2 concentration - sensor interface | Maxim | MAX232CPE+ | Type: Multichannel RS-232 driver | S8 | n/a | U1 |
| Sensing | Air CO2 concentration - capacitor | Tru Components | 442896 | Capacitance: 1 µF | S8 | n/a | C1-C5 |
| Sensing | Air flow rate - sensor module | Honeywell | HAFUHT0100L4AXT | Range: 0…1000 L/min | 1, S4, S8 | n/a | J3 |
| Sensing | Air pressure - sensor module | WIKA | A-10-6-BA316-HD1Z-AA-M4Z-ZW | Range: 0...1.6 bar | 1, S4, S8 | n/a | J1 |
| Sensing | Air pressure - resistor | TE Connectivity | YR1B165RCC | Resistance: 165 Ohm, Power: 0.25 W | S8 | n/a | R1 |
| Sensing | Air relative humidity - sensor module | TE Connectivity | HM1500LF | Range: 0...100 % | 1, S4, S8 | n/a | J2 |
| Sensing | Air relative humidity - operation amplifier | Texas Instruments | OPA340PA | Output Current per Channel: 50 mA | S8 | n/a | U2 |

| | | | | | | | |
|---|---|---|---|---|---|---|---|
| Sensing | Air relative humidity - operation amplifier | Texas Instruments | LM358APE4 | Output Current per Channel: 20 mA | S8 | n/a | U1 |
| Sensing | Air relative humidity - resistor | Vishay | MRS16000C2002FCT00 | Resistance: 20 kOhm, Power: 0.4W, Accuracy: ±1% | S8 | n/a | R2, R3 |
| Sensing | Air relative humidity - resistor | Vishay | MRS16000C1201FCT00 | Resistance: 1.2 kOhm, Power: 0.4W, Accuracy: ±1% | S8 | n/a | R1, R4 |
| Sensing | Air temperature - sensor module | WIKA | TF35 | Range: -50...+250 °C | 1, S4, S8 | n/a | TH4 |
| Sensing | Air temperature - instrument amplifier | Texas Instruments | In/a125PA | Mounting: THT | S8 | n/a | IC1 |
| Sensing | Air temperature - resistor | Arcol | MRA020710KBTA015 | Resistance: 10 kOhm, Power: 0.25W | S8 | n/a | R5 |
| Sensing | Air temperature - bridge resistors | VPG Foil Resistors | Y00621K00000T9L | Resistance: 1 kOhm, Power: 0.25 W, Accuracy: ±0.01% | S8 | n/a | R1, R2, R3 |
| Sensing | Air temperature - capacitor | KEMET | Goldmax 300 MLCC | Capacitance: 0.1 µF | S8 | n/a | C1-C5 |
| Actuation | Air circulation - pump | Boxer Pumps | 3112.660 | Type: Brushless, diaphragm | 1,S4, S7 | P1-P7 | n/a |
| Actuation | Air circulation - power supply | TT Electronics | SRS150-24 | Output voltage: 24 V | S7 | n/a | n/a |
| Actuation | Air circulation - potentiometer | Bourns | 3296W-1-102LF | Max. resistance: 1 kOhm, Accuracy: ±10% | S7 | Pt1-Pt7 | n/a |
| Actuation | Air circulation - voltage regulator | STMicroelectronics | L7805A | Output voltage: 5 V | S7 | n/a | n/a |
| Actuation | Air circulation - printer circuit board | Multi Leiterplatten GmbH | n/a | Material: FR4 1.55mm, Copper plane: 35 µm | S7 | n/a | n/a |
| Actuation | Air switching - valve | SMC | VDW350-5G-4-02F-Q | Type: 3/2-way | 1, S4, S7 | UV1-UV6, DV1-DV6, AV | n/a |
| Actuation | Air switching - relais | Pan/asonic | AQY211EH | Type: solid state relais | S8 | n/a | K1 |
| Actuation | Air switching - diode | Diotec | 1N4001 | Nomin/al current: 1A | S8 | n/a | D |
| Actuation | Water circulation pump | TSC | MGD1000S | Type: Brushless, geared | 1, S5, S6 | WP0 | n/a |
| Actuation | Water circulation control board | TSC | EQi-UART-MG1 | Type: Brushless | S6 | n/a | n/a |
| Actuation | Water switching valve | SMC | VDW22LA | Type: 2/2-way | S5, S6 | IV1-IV6 | n/a |
| Actuation | Water switching valve adapter | SMC | KQ2H04-M5A | Material: brass | n/a | n/a | n/a |
| Actuation | Water switching push button | Würth | 430156050726 | Type: SPST | S6 | PB1-PB6 | n/a |
| Actuation | Water switching relais | Pan/asonic | AQY211EH | Type: solid state relais | n/a | n/a | n/a |
| Media handling | Air filter (centrifgual/sintering) | Landefeld | EAF 2000-02 | Membrane: Pore size: 5 µm | S4 | n/a | n/a |
| Media handling | Air filter (hydrophobic) | Sartorius | Midisart 2000 | Membrane: PTFE, Pore size: 0.2 µm | 1, S4, S5 | n/a | n/a |
| Media handling | Air perforated tubes | Landefeld | PUN 6X4 KLAR | Material: PU, Shore hardness:95 A | n/a | n/a | n/a |

| | | | | | | | |
|---|---|---|---|---|---|---|---|
| Media handling | Water manifold (extern/al) | SMC | KM11-04-08-6 | Material: PBT | S5 | n/a | n/a |
| Media handling | Water supply tube | Landefeld | PUN 4X2,5 KLAR | Material: PU, Shore hardness:95 A | n/a | n/a | n/a |
| Media handling | Water non-return valve | Bürkle | 229-0134 | Opening pressure: 0.86 mbar | S5 | n/a | n/a |
| Media handling | Water filter (hydrophilic) | Sartorius | Sartolab P20 | Membrane: PES, Pore size: 0.2 µm | 1, S5 | n/a | n/a |
| Media handling | Water supply tube | Landefeld | PUN 8X5 KLAR | Material: PU, Shore hardness:95 A | n/a | n/a | n/a |
| Media handling | Water vessel | Schott | GL45 | n/a | 1, S5 | n/a | n/a |
| Media handling | Water straight adapter (at EcoChamber) | SMC | KQ2N06-99 | Material: PBT | n/a | n/a | n/a |
| Media handling | Water distribution port (at manifold) | Qosin/a | 11548 | Material: polycarbon/ate | n/a | n/a | n/a |
| Media handling | Water manifold (intern/al) | Formlabs | n.a. (custom) | Material: Biomed Resin (amber) | n/a | n/a | n/a |
| Media handling | Water distribution port (to metal tray) | Qosin/a | 11544 | Material: polycarbon/ate | n/a | n/a | n/a |
| Media handling | Water distribution tube (to metal tray) | Landefeld | SILI 1,52,5 | Material: silicone | n/a | n/a | n/a |
| Media handling | Water distribution port (to flow pots) | Qosin/a | 11545 | Material: polycarbon/ate | n/a | n/a | n/a |
| $CO_2$ atmosphere | Air compressor for CO2 scrubber | Clean/air | CLR 20/50 T | Max. pressure: 10 bar. Vessel volume: 50 L | 1 | n/a | n/a |
| $CO_2$ atmosphere | Purge gas generator (CO2 scrubber) | CMC Instruments | PG28L | Membrane: Pore size: 0.1 µm | 1 | n/a | n/a |
| $CO_2$ atmosphere | Over-pressure relief valve | Landefeld | SV9 2055 7.5 D/G 0,48 | Max. set pressure: 0.4 bar | n/a | n/a | n/a |
| $CO_2$ atmosphere | Butterfly valve | Landefeld | KH 14 DVGW KN | Pressure range: -0.9…+50 bar | n/a | n/a | n/a |
| $CO_2$ atmosphere | Pressure controller | Landefeld | R 014-1,5 MS-CO | Pressure range: 0.1…+1.5 bar | n/a | n/a | n/a |
| $CO_2$ atmosphere | Switch valve | GOK | AUV PS 16 | Max. pressure: 16 bar | n/a | n/a | n/a |
| $CO_2$ atmosphere | Pressure reducer | Air Liquide | Primejet 200-10-30 | Max output pressure: 10 bar | n/a | n/a | n/a |
| $CO_2$ atmosphere | Carbon dioxide cylinder | Air Liquide | I6510L50R0A001 | Volume: 50 L, Type: LASAL 2 | 1 | n/a | n/a |
| $CO_2$ atmosphere | Tube for extern/al air supply | Festo | PUN-H-6X1-TGE TPU | Max pressure: 10 bar | 1 | n/a | n/a |
| Auxiliary components | Air pump service kit | Boxer Pumps | 3000.137 | n/a | n/a | n/a | n/a |
